# Selection on old variants drives adaptive radiation of *Metrosideros* across the Hawaiian Islands

**DOI:** 10.1101/2020.11.03.354068

**Authors:** Jae Young Choi, Xiaoguang Dai, Julie Z. Peng, Priyesh Rughani, Scott Hickey, Eoghan Harrington, Sissel Juul, Julien Ayroles, Michael Purugganan, Elizabeth A. Stacy

## Abstract

Some of the most spectacular adaptive radiations begin with founder populations on remote islands. How genetically limited founder populations give rise to the striking phenotypic and ecological diversity characteristic of adaptive radiations is a paradox of evolutionary biology. We conducted an evolutionary genomic analysis of genus *Metrosideros*, a landscape-dominant, incipient adaptive radiation of woody plants that spans a striking range of phenotypes and environments across the Hawaiian Islands. Using nanopore-sequencing, we created a chromosome-level genome assembly for *M. polymorpha* var. *incana* and analyzed wholegenome sequences of 131 individuals from 11 taxa sampled across the islands. We found evidence of population structure that grouped taxa by island. Demographic modeling showed concordance between the divergence times of island-specific lineages and the geological formation of individual islands. Gene flow was also detected within and between island taxa, suggesting a complex reticulated evolutionary history. We investigated genomic regions with increased differentiation as these regions may harbor variants involved in local adaptation or reproductive isolation, thus forming the genomic basis of adaptive radiation. We discovered differentiation outliers have arisen from balancing selection on ancient divergent haplotypes that formed before the initial colonization of the archipelago. These regions experienced recurrent divergent selection as lineages colonized and diversified on new islands, and hybridization likely facilitated the transfer of these ancient variants between taxa. Balancing selection on multiple ancient haplotypes–or time-tested variants–may help to explain how lineages with limited gene pools can rapidly diversify to fill myriad ecological niches on remote islands.

**Significance statement:** Some of the most spectacular adaptive radiations of plants and animals occur on remote oceanic islands, yet such radiations are preceded by founding events that severely limit genetic variation. How genetically depauperate founder populations give rise to the spectacular phenotypic and ecological diversity characteristic of island adaptive radiations is not known. We generated novel genomic resources for Hawaiian *Metrosideros*––a hyper-variable incipient adaptive radiation of woody taxa—for insights into the paradox of remote island radiations. We found that *Metrosideros* colonized each island shortly after formation and diversified within islands through recurrent selection on ancient variations that predate the radiation. Recurring use of ancient variants may explain how genetically depauperate lineages can diversify to fill countless niches on remote islands.

## Introduction

Adaptive radiations exhibit extraordinary levels of morphological and ecological diversity (1). Although definitions of adaptive radiation vary (2–7), all center on ecological opportunity as a driver of adaptation and, ultimately, diversification (2, 8–10). Divergent selection, the primary mechanism underlying adaptive radiations, favors extreme phenotypes (11) and selects alleles that confer adaptation to unoccupied or under-utilized ecological niches. Differential adaptation results in divergence and, ultimately, reproductive isolation between populations (12). Adaptive radiations demonstrate the remarkable power of natural selection as a driver of biological diversity, and provide excellent systems for studying evolutionary processes involved in diversification and speciation (13).

Adaptive radiations on remote oceanic islands are especially interesting, as colonization of remote islands is expected to involve population bottlenecks that restrict genetic variation (14). Adaptive radiations in such settings are especially impressive and even paradoxical, given the generation of high species richness from an initially limited gene pool (15). Several classic examples of adaptive radiation occur on oceanic islands, such as Darwin’s finches from the Galapagos islands (16), anole lizards from the Caribbean islands (9), Hawaiian Drosophilids (17), and Hawaiian silverswords (18), to name a few.

Recent advances in genome sequencing and analyses have greatly improved our ability to examine the genetics of speciation and adaptive radiations. By examining sequences of multiple individuals from their natural environment, it has become possible to “catch in the act” the speciation processes between incipient lineages (19). Genomic studies of early-stage speciation show that differentiation accumulates in genomic regions that restrict the homogenizing effects of gene flow between incipient species (20). The number, size, and distribution of these genomic regions can shed light on evolutionary factors involved in speciation (19). Regions of high genomic differentiation can also form from evolutionary factors unrelated to speciation, such as linkage associated with recurrent background selection or selective sweeps on shared genomic features (21, 22).

Genomic studies of lineages undergoing rapid ecological diversification have begun to reveal the evolutionary mechanisms underlying adaptive radiations. Importantly, these studies highlight the pivotal role of hybridization between populations and the consequent exchange of adaptive alleles that facilitates rapid speciation and the colonization of diverse niches (23–25). Most genomic studies of adaptive radiation involve animal systems, however, in particular, birds and fishes. In plants, genomic studies of adaptive radiations are sparse (26–28) and all examine continent-wide radiations, while there are no equivalent genomic studies of plant adaptive radiations in geographically restricted systems such as remote islands. Because the eco-evolutionary scenarios associated with adaptive radiations are diverse (5, 29), whether commonalities identified in adaptive radiations in animals (23, 30) are applicable to plants is an open question. For example, the genetic architecture of animal adaptive radiations typically involves differentiation at a small number of genomic regions (31–33). In contrast, the limited insights available for plants suggest a more complex genetic architecture (26).

We investigated the evolutionary genomics of adaptive radiation in *Metrosideros* Banks ex Gaertn. (Myrtaceae) across the Hawaiian archipelago. Hawaiian *Metrosideros* is a landscape-dominant, hypervariable and highly dispersible group of long-lived [possibly > 650 years] (34) woody taxa that are non-randomly distributed across Hawaii’s heterogeneous landscape, including cooled lava flows, wet forests and bogs, subalpine zones, and riparian zones (35, 36). About 25 taxa or morphotypes are distinguished by vegetative characters ranging from prostate plants that flower a few centimeters above ground to 30-m tall trees, and leaves range dramatically in size, shape, pubescence, color, and rugosity (35, 37, 38). Variation in leaf mass per area within the four *Metrosideros* taxa on Hawaii Island alone matches that observed for woody species globally (39). Common garden experiments (38, 40–44) and parent-offspring analysis (45) demonstrate heritability of taxon-diagnostic vegetative traits, indicating that taxa are distinct genetic groups and not the result of phenotypic plasticity. *Metrosideros* taxa display evidence of local adaptation to contrasting environments (46, 47), suggesting ecological divergent selection is responsible for diversification within the group (48). This diversification, which spans the past ~3.1-3.9 million years (MY) (49, 50), has occurred in spite of the group’s high capacity for gene flow by way of showy bird-pollinated flowers and tiny wind-dispersed seeds (36, 51). Lastly, the presence of partial reproductive isolating barriers between taxa is consistent with the early stages of speciation (52). Here, we generated several genomic resources for Hawaiian *Metrosideros* and used these in population genomic analyses to gain deeper insights into the genomic architecture and evolutionary processes underlying this island adaptive radiation.

## Results

### An improved genome assembly for the Hawaiian *Metrosideros*

Using nanopore sequencing, an individual *Metrosideros polymorpha* var. *incana* was sequenced to 66× coverage (see Table S1 for genome sequencing statistics). The reads were assembled into a draft assembly that had high contiguity with a contig N50 of 1.85 Mbp (Table 1). We implemented Pore-C sequencing (53), which combines chromosome conformation capture with long-read nanopore sequencing, to assay the *Metrosideros*-specific chromosome contact map and anchor contigs to their chromosomal positions [see Table S2 for Pore-C sequencing statistics] (54). Using Pore-C contact maps, initial assembly contigs were scaffolded into 11 super-scaffolds (Fig. 1A) spanning 292.8 Mbps with an N50 of 25.9 Mbp. Compared to a previous genome assembly that was based only on Illumina sequencing (55), our assembly was similar in total genome size but had significantly higher contiguity. The number of super-scaffolds was consistent with the 11 chromosomes in *Metrosideros* (56). The assembly was evaluated with 2,326 Benchmarking Universal Single-Copy Ortholog (BUSCO) genes from eudicots, and 2,183 genes (93.8%) were present. These results suggest that our chromosome-level genome assembly was highly contiguous and complete. Gene annotation was conducted using nanopore sequencing of a cDNA library generated from leaf tissue (see Table S3 for cDNA sequencing statistics). A total of 28,270 genes were predicted with 94.2% of the transcripts showing an annotation edit distance (AED) of less than 0.5.

**Figure 1.**
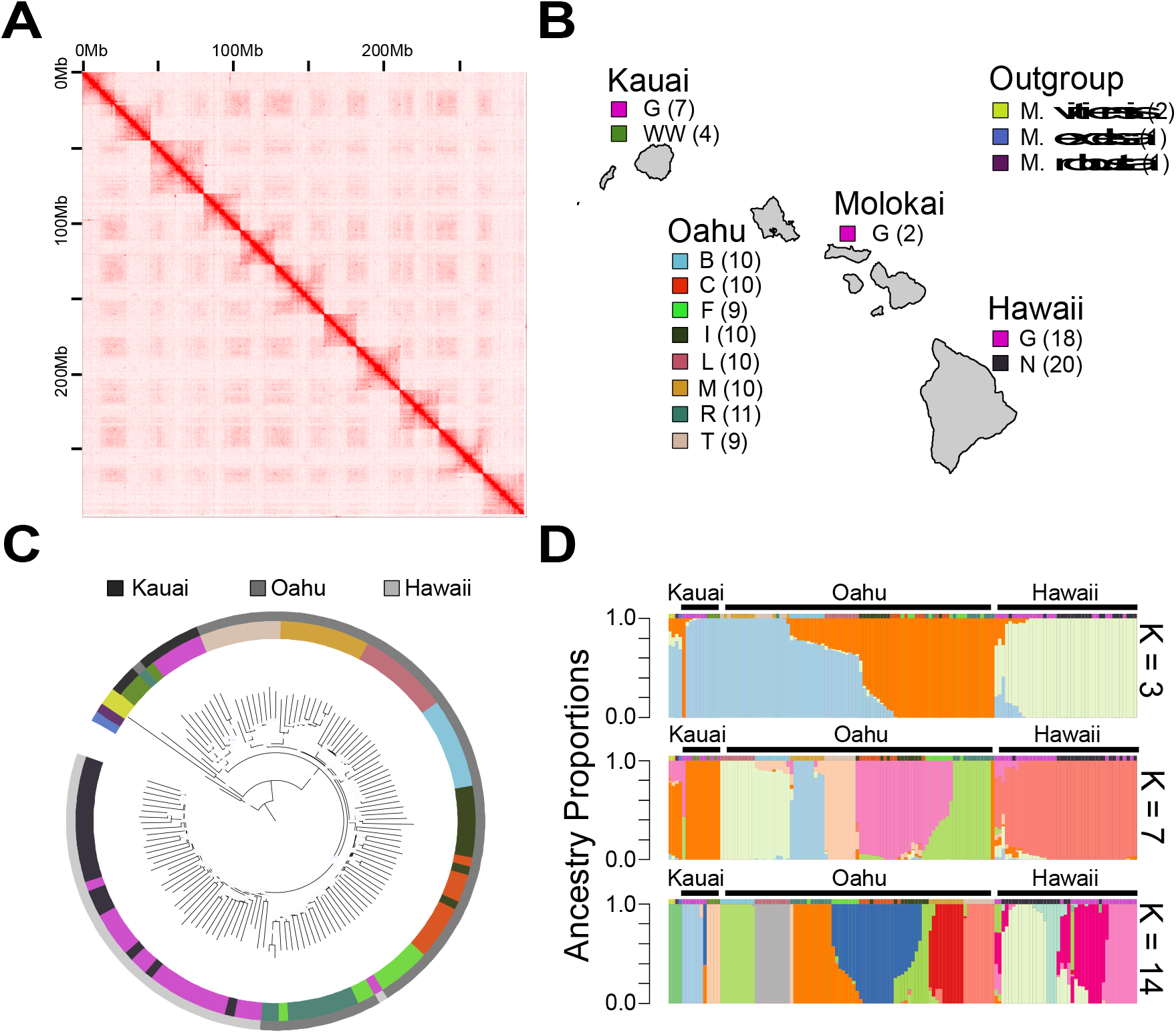
Genomics of Hawaiian *Metrosideros*. (A) Pore-C-based chromosome contact matrix for *M. polymorpha* var. *incana*. (B) Geographic distribution and taxon classification for the 135 samples that were analyzed in this study. Numbers in parentheses represent sample sizes. (C) Genome-wide neighbor-joining tree built using a distance matrix. Outer circle colors indicate island of origin for each sample, and inner circle colors indicate taxa as in panel (B). Nodes with greater than 95% bootstrap support are indicated with light blue circles. (D) Ancestry proportion estimates using the ADMIXTURE algorithm for K = 3, 7, and 14. Colors above admixture barplots represent taxa as in panel (B).

**Table 1.** Genome assembly statistics for *M. polymorpha* var. *incana*.

| Assembly features |  |
| --- | --- |
| # of contigs | 1,035 |
| # of scaffolded psuedomolecules | 11 |
| Total number of bases in contigs | 307,350,354 bp |
| Total number of bases scaffolded | 292,766,828 bp |
| Maximum contig length | 8,272,703 bp |
| Contig N50 length | 1.852 Mbp |
| Scaffold N50 length | 25.93 Mbp |
| BUSCO score | 93.8% |
| GC content | 39.8% |
| Repeat content | 36.3% |
| Number of annotated genes | 28,270 |

The population genomics of Hawaiian *Metrosideros* was investigated by whole-genome re-sequencing 89 individuals from the islands of Oahu and Kauai, and combining these data with previously published sequence data from Hawaii Island and Molokai (57). In addition, we sequenced 4 *Metrosideros* species outside the Hawaiian archipelago as outgroup genomes. In total, we analyzed 131 individuals belonging to 11 taxa (Taxa are abbreviated as *M. polymorpha* race B [B], *M. polymorpha* race C [C], *M. polymorpha* race F [F], *M. polymorpha* race L [L], *M. polymorpha* var. *glaberrima* [G], *M. polymorpha* var. *incana* [I], *M. macropus* [M], *M. polymorpha* var. *newellii* [N], *M. polymorpha* var. *waialealae* [WW], *M. rugosa* [R], and *M. tremuloides* [T]) across the Hawaiian archipelago (Fig 1B). The median genome coverage was 14× per individual, and on average 93% of the sequencing reads were aligned to our reference genome (Table S4). The mapped population genomic data were used to call SNPs and after filtering there were 22,511,205 variable sites that were used for subsequent analysis.

### Population structure and relationships across the Hawaiian archipelago

A neighbor-joining phylogenomic tree was reconstructed to examine evolutionary relationships among populations (Fig. 2C). The tree topology grouped individuals by island with little evidence of recent migration between islands. Within the phylogeny, individuals clustered according to taxonomic designations and were monophyletic with high confidence (>95% bootstrap support). Exceptions were paraphyletic relationships among taxa C, F, and I on Oahu and taxa G and N on Hawaii Island. Among the outgroup species, *M. vitiensis* from Fiji and American Samoa was sister to all Hawaiian *Metrosideros*.

**Figure 2.**
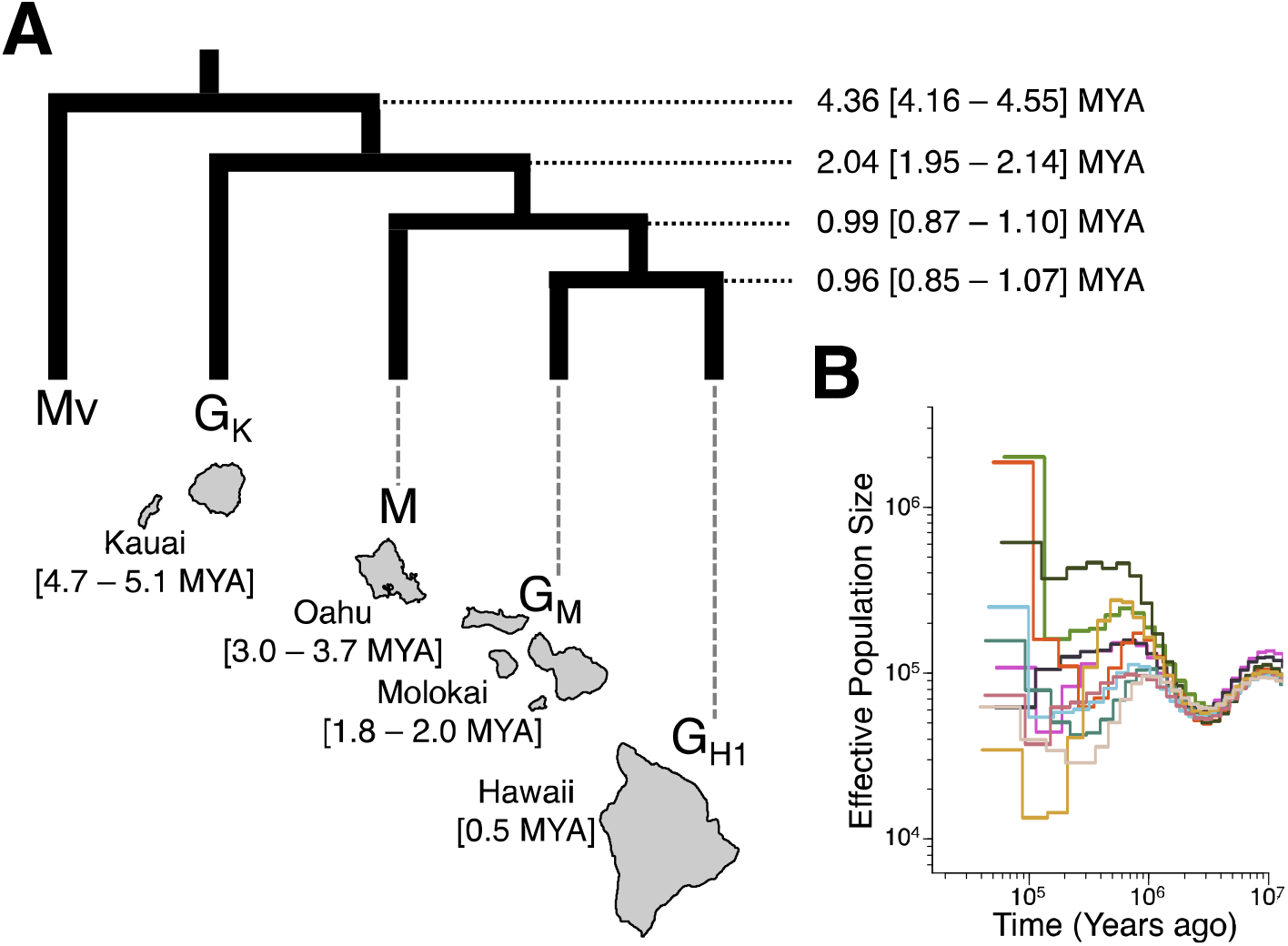
Divergence time and demographic history of Hawaiian *Metrosideros*. Relative times were converted to absolute times assuming a mutation rate of 7e^−9^ mutations per base pair per generation and a 20-year generation time. (A) G-PhoCS-estimated divergence times for representative taxa from each island (above), and time of geological formation of each island based on Clague (62) (below). (B) MSMC2-estimated effective population size for each Hawaiian taxon, color-coded as in Fig. 1B.

Finer scale evolutionary relationships across Hawaiian *Metrosideros* were examined by estimating genomic ancestry proportions (K) for each individual (see Fig. S1 for K = 3 to K = 15 results). At K=3 most individuals from Kauai and Hawaii Island showed a single, island-specific ancestry component (Fig 2D). On Oahu, there were taxa with Oahu-specific ancestry (taxa C, F, and R), Kauai-specific ancestry (taxa M and T), and mixed Oahu and Kauai ancestry (taxa B, L, and I). With increasing K each taxon/population showed increasingly unique ancestry, and at K = 14, with few exceptions, individuals belonging to the same taxon/population shared a single, unique ancestry. One exception was taxon F, for which all individuals consistently showed admixed ancestry comprising components of both taxa C and R. Additionally, for taxa G and N on Hawaii Island, increasing K led to further subdivision of ancestry, consistent with the complex population substructure of Hawaii Island *Metrosideros* (57). On Hawaii Island, G belonged to two genetic groups designated G_H1_ and G_H2_ in our previous analysis, and G_H2_ represented a recently admixed population with taxon N (57). Taxon F on Oahu and population G_H2_ on Hawaii Island are likely to be hybrid populations formed from recent hybridization of genetically distinct populations (58). As such, they were removed from downstream analysis. We focused on 10 taxa/populations (B, C, G_H1_, G from Kauai [G_K_], I, L, M, N, R, and T), each of which had a largely single ancestry and sufficient sample sizes (minimum of 7 individuals in G_K_ and maximum of 14 individuals in taxon N) for population genomic analysis.

We investigated evidence for gene flow between taxa by calculating Patterson’s D-statistics (ABBA-BABA D test) (59, 60) on all population trios following the species-wide topology (Fig. 2C). *M. vitiensis* was used as the outgroup, specifically the sample from Fiji due to its high genome coverage. Overall, 85 of the 159 (53.5%) trio topologies had significant D-statistics (Bonferroni corrected p-value < 0.05; see Fig. S2 for results of each trio). Of the 85 trios, 53 (62.4%) topologies involved admixture between taxa on different islands, indicating that reticulate evolution was pervasive within and between islands. Taxa M and T, however, were exceptions, as they had the fewest significant D-statistics and no evidence of admixture with lineages outside of Oahu (Fig. S2).

### Divergence times of *Metrosideros* across the islands

We investigated the speciation history of Hawaiian *Metrosideros* through demographic modeling. The population divergence times were estimated using the generalized phylogenetic coalescent sampler (G-PhoCS) (61). From each island, a single population/taxon was chosen as a representative since we were interested in the colonization history of *Metrosideros* across the islands. Populations of the archipelago-wide taxon G were chosen from each island, except Oahu, for which G samples were not available. Thus, taxon M was chosen to represent Oahu, as it had no significant evidence of between-island admixture (Fig. S2), which reduced the number of admixture models to be tested with G-PhoCS.

Initially, we ran G-PhoCS models fitting migration bands (*i.e*. admixture) between terminal lineages. Results showed that parameterizing admixture had no effect on estimates of divergence times (Fig. S3). Given this result, we based our divergence time analysis on the simple no-migration model. The G-PhoCS-estimated divergence time between Hawaiian *Metrosideros* and the outgroup *M. vitiensis* was 4.36 million years ago [MYA] (95% Highest Posterior Density [HPD] 4.16 – 4.55 MYA), which is younger than the geological formation of Kauai (Fig. 2A). However, given that *M. vitiensis* is not the most closely related outgroup species to Hawaiian *Metrosideros* (50), colonization of the Hawaiian Islands likely occurred more recently than 4.36 million years ago. Additionally, the divergence time estimates within the Hawaiian archipelago suggested that colonization of each island occurred long after its formation. The exception was Hawaii Island for which the divergence time of *Metrosideros* predated the geological formation of that island, consistent with the results of our previous study (57).

We used the program Multiple Sequentially Markovian Coalescent 2 [MSMC2] (63, 64) to estimate past changes in effective population size (N_e_) for each taxon/population. Results showed that all taxa had identical trajectories in showing a decrease in N_e_ until ~3 MYA, followed by an increase and subsequent drop in N_e_ in a pattern unique to each taxon (Fig. 2B). This result suggests that all Hawaiian *Metrosideros* taxa share the same common ancestor that experienced a population bottleneck about 3 – 4 MYA when the ancestral population initially colonized the archipelago. Based on G-PhoCS and MSMC2 analysis the initial colonization of the Hawaiian archipelago was estimated to have occurred ~3 – 4.4 MYA (150,000–220,000 generations ago).

### The genomic landscape of differentiation

To investigate the genetic architecture underlying the *Metrosideros* radiation we narrowed our analysis to pairs of taxa that were phylogenetic sisters, since for these sister pairs the pattern of genome-wide differentiation reflects the early stages of speciation/radiation. For these sisters we used δaδi (65) to fit 20 different demography models (see Table S5 for complete δaδi results) to find the best-fitting model. Results showed that pair G_H1_ and N and pair C and I were consistent with a speciation model in which lineage divergence occurred with continuous gene flow [*i.e*. primary gene flow] (66), while in pair B and L and pair M and T the populations have been largely isolated from each other with the exception of either a recent or ancient gene flow event (Fig 3A).

**Figure 3.**
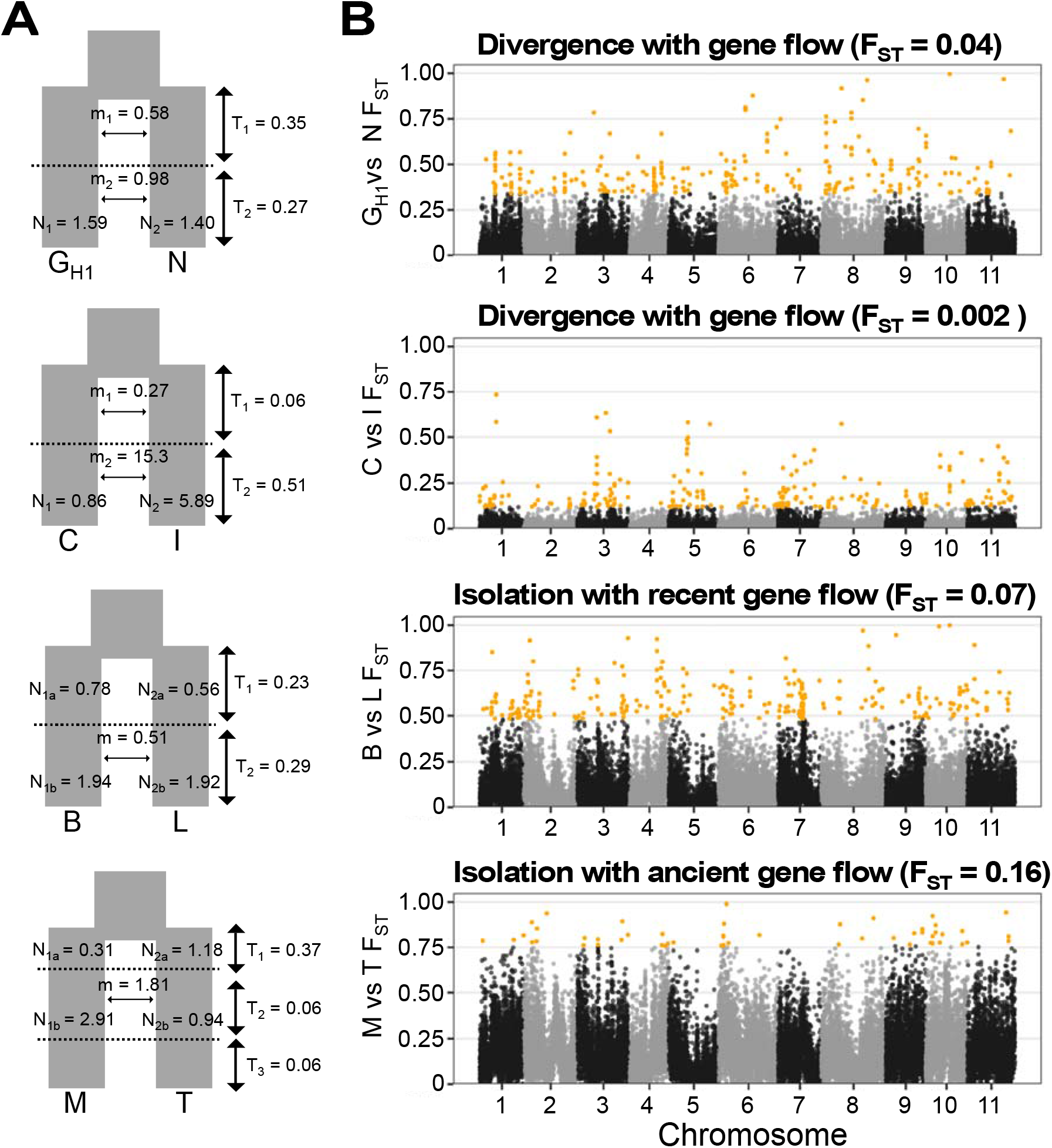
Genomic landscape of differentiation for the four phylogenetic sister pairs. (A) bestfitting demography model based on δaδi modeling. (B) Genome-wide F_ST_ in 10-kbp windows. Yellow dots are outliers identified with z-score-transformed F_ST_ values (zF_ST_) > 4.

We investigated the genomic architecture of the *Metrosideros* adaptive radiation by quantifying genome-wide patterns of differentiation and signatures of divergent selection between sister taxa. We focused on differentiation (F_ST_) outlier regions since these regions would harbor genetic variation associated with the divergence of sister pairs (67). Results showed that, in all four sister pairs, areas of high genomic differentiation were scattered across all 11 chromosomes (Fig. 3B). Pair M and T had the fewest outlier windows (52 zF_ST_ outlier windows), while taxon pairs G_H1_ and N, C and I, and B and L had over 200 zF_ST_ outlier windows each (257, 226, and 260, respectively). The median genome-wide F_ST_ between M and T (F_ST_ = 0.16) was more than twice the level of F_ST_ within other sister pairs (G_H1_ and N median F_ST_ = 0.04; C and I median F_ST_ = 0.002; B and L median F_ST_ = 0.07), suggesting that increased genome-wide differentiation between M and T due to their genetic isolation may have eroded outlier windows to undetectable levels (26, 68). The genomic positions of outlier windows generally did not overlap across four sister-taxon pairs (*i.e*. >84% of outlier windows were found in only one taxon pair; see Fig. S4).

### Evolutionary factors shaping genomic differentiation

Genomic outliers of differentiation do not always result from divergent selection (69–71), and F_ST_ can be a biased estimate of differentiation (72). Hence, as a complementary analysis we examined levels of absolute genetic divergence (D_xy_) within differentiation outlier regions, as D_xy_ is expected to be elevated in regions under divergent selection or regions acting as genetic barriers between populations (69). Within each pair of sister taxa, D_xy_ levels were significantly elevated in differentiation outliers (MWU test p-value < 0.001) relative to genomic background (Fig. 4A left box). Further, genomic differentiation outlier regions had significantly elevated D_xy_ levels when compared with other non-sister taxa representing increasing phylogenetic distances [MWU test p-value < 0.05] (Fig. 4A right box). Interestingly, differentiation outliers had elevated D_xy_ even in comparisons with G_K_ on the oldest island, Kauai.

**Figure 4.**
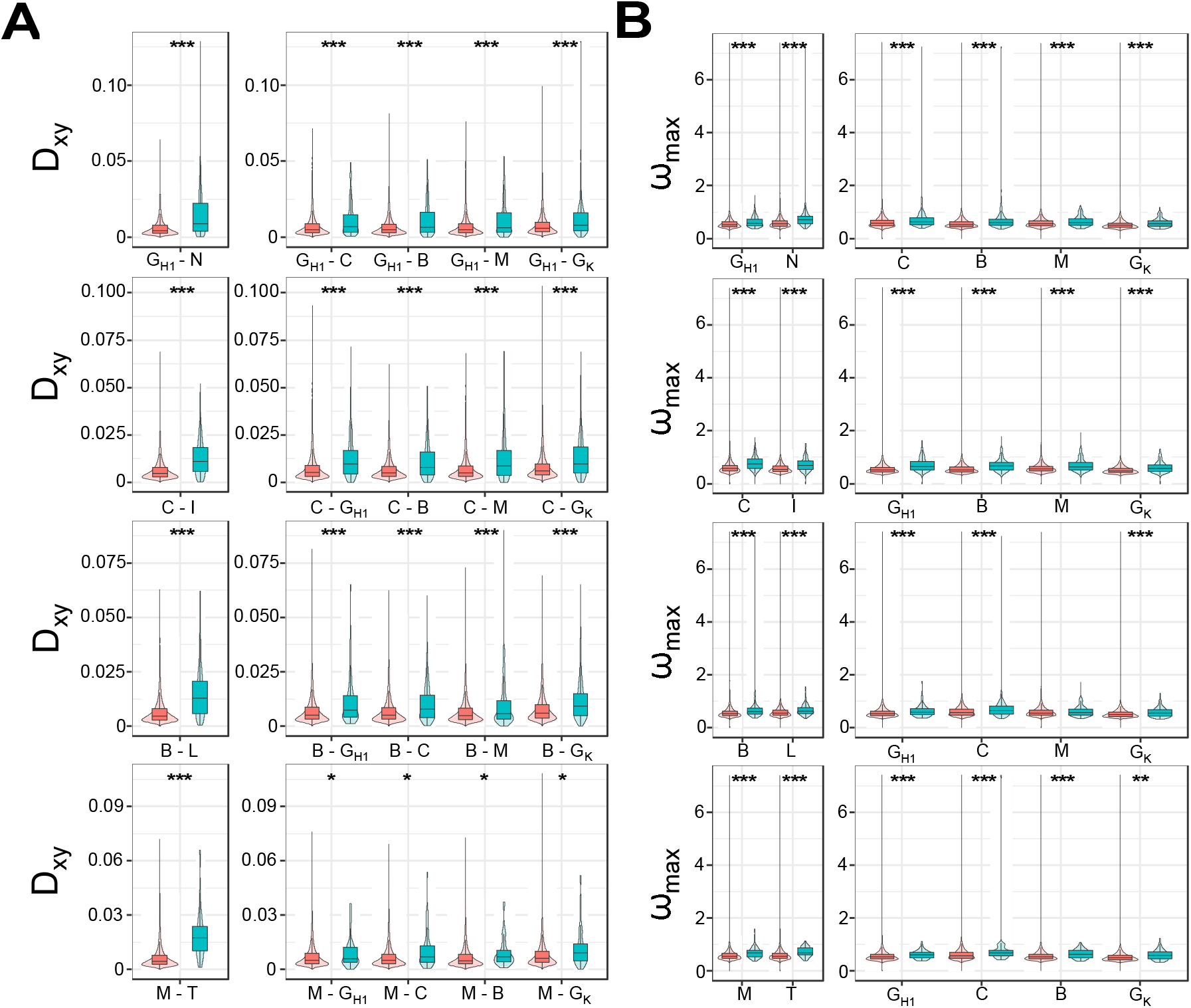
Sequence divergence and evidence of selective sweeps in differentiation outlier regions. Red boxes are statistics from the genomic background, and green boxes are statistics from the differentiation outlier regions. (A) Sequence divergence (D_xy_) statistics identified in a sister pair (left window) and D_xy_ estimated from the same regions but between one member of the pair and four other taxa representing increasing phylogenetic distances (right window). (B) Selective sweep statistics (ω_max_) identified in a sister pair (left window) and the same regions in other *Metrosideros* taxa (right window). * indicate p < 0.05, ** indicate p < 0.01, and *** indicate p < 0.001.

We examined evidence of selective sweeps in regions of elevated genomic differentiation using ω_max_ statistics. Results showed ω_max_ were significantly higher for differentiation outlier regions (MWU test p-value < 0.001) compared to genomic background (Fig. 4B left box). Further, the same regions had significantly elevated ω_max_ (MWU test p-value < 0.01) in other taxa as well (Fig. 4B right box), indicating that the differentiation outlier regions formed between sister pairs were also targets of positive selection in multiple other *Metrosideros* taxa.

These results suggest that an archipelago-wide selective pressure has acted recurrently on the same genomic regions across multiple *Metrosideros* taxa. Such recurrent selection can have profound effects on the genomic landscape of differentiation between populations (27, 73). In *Metrosideros* we found that π, D_xy_, F_ST_, and ω_max_ were all positively and significantly correlated among taxa (Fig. S5). Because archipelago-wide selection is unlikely to be involved in the early-stage divergence of lineages, we examined regions that had evidence of elevated genomic differentiation in specific lineages. Using the correlated differentiation landscape as a genomic control (21), we focused on regions that were significantly differentiated in individual sister pairs (see Table S6 for genomic positions). D_xy_ within these lineage-specific differentiation regions were significantly elevated compared to the genomic background (MWU test p-value < 0.001) [Fig. S6]. When compared to other non-sister taxa with increasing phylogenetic distances, the lineage-specific differentiation regions still had elevated D_xy_ levels compared to genomic background (MWU test p-value < 0.01), except for the pair M and T.

A gene ontology-(GO) enrichment analysis was conducted to identify any over-represented functional categories associated with regions of genomic differentiation. Results showed that genes within the outlier windows were enriched for 19 GO terms in the ‘biological process’ category, and the functions were largely related to metabolic processes, cell cycle, and disease immunity responses (Fig. S7).

## Discussion

The evolution of plant adaptive radiations on islands has been studied largely through phylogenetic approaches by comparing DNA sequences or genomes of a single representative from each species within the radiation (74, 75). In contrast, we investigated the evolution of *Metrosideros* by constructing a chromosome-level genome assembly and sampling genome-wide variation of 131 individuals across the Hawaiian Islands. Using a population genomics approach, we inferred the evolutionary history of Hawaiian *Metrosideros*, including population divergence times and demography associated with this island adaptive radiation. We discovered that selection on ancient variation played a major role in shaping regions of genome-wide differentiation during the recent and ancient radiation of this group across the Hawaiian Islands.

The timing of colonization of the Hawaiian Islands by *Metrosideros* may explain the contrast in current *Metrosideros* diversity and dominance on the two oldest main islands. Coalescent analysis yielded an estimate for the initial split between the common ancestor of all Hawaiian *Metrosideros* and the outgroup *M. vitiensis* at 4.4 MYA, when only Kauai, the oldest main island, was present for colonization, having completed the primary (shield-building) phase of island growth (62, 76). Phylogeography and population structure analysis indicated that the radiation was endogenous to the islands wherein colonization and subsequent lineage diversification occurred within newly formed islands in the order of their appearance. This progressive colonization from older to younger islands [*i.e*. progression rule] (77) is a hallmark phylogeographic pattern for many Hawaiian lineages (78). Our lower bound estimate of the initial colonization of the islands, however, was 3 MYA, consistent with the estimate (3.1 MYA) derived from a recent fossil-calibrated analysis of 40 nuclear genes (50). This estimate of a more recent colonization of the islands is consistent with the lower dominance of *Metrosideros* on Kauai relative to the younger islands and the greater taxonomic richness of *Metrosideros* on islands of intermediate age, especially Oahu (35). Both of these patterns would be expected if *Metrosideros* colonized the islands more recently when Oahu was nascent and Kauai was fully formed and already colonized by lineages from the northwest Hawaiian Islands or more distant sources (*e.g*., lobelioids (79), silverswords (80), *Psychotria* (81)). With the available data, it is not possible to determine the order or exact timing of island colonization for *Metrosideros*. Increased genomic sampling within the south Pacific island chains and along the hypothesized origins of the Hawaiian lineage (49, 50) could clarify the colonization history and ancestral diversity of this group.

The phylogenetic sister taxa that we examined inhabit contrasting environments [G_H1_ vs. N: wet forest vs. riparian; C vs. I: lower montane vs. low-elevation, dry, including new lava flows; B vs. L: cloud forest vs. upper montane; M vs. T: montane vs. steep windward slope] (35, 36), and differential local adaptation has been suggested as the primary driver of their phenotypic divergence (46, 47). The genome-wide distribution of differentiation outliers suggests that ecological diversification within *Metrosideros* either has a polygenic basis or involves multiple traits with simple genetic architectures. *Metrosideros* taxa display a rich diversity of vegetative traits that are expected to be polygenic (82, 83), consistent with the differentiation outliers that were well distributed throughout the eleven chromosomes. The number of differentiated genomic regions was not associated with migration rate between sister taxa, suggesting that differentiation outliers were not necessarily formed as barriers to gene flow (20). These patterns contrast with the genetic architecture observed for key traits in animal adaptive radiations. For instance, in butterflies (84), crows (85), and cichlids (86) where adaptive radiation involves different color morphs, one finds a small number of genomic regions that are strongly differentiated due to the simple genetic architecture of color-based traits, which also contribute to assortative mating and further build up differentiation (87). In animal adaptive radiations, in general, differentiation is seemingly localized to a few genomic regions with prominent broad peaks (31–33). In contrast, the outlier peaks in *Metrosideros* were narrow and their distribution was heterogeneous across the genome, a pattern that was also found in the continent-wide adaptive radiation of sunflowers (26). The narrow peaks suggest that fine-scale mapping of the genes underlying divergent phenotypes may be possible.

DNA divergence and selective sweep analyses of the differentiation outliers suggest a prominent role for ancient balanced polymorphisms in the radiation of *Metrosideros* in Hawaii. The differentiation outliers in *Metrosideros* showed evidence of selective sweeps in multiple taxa, suggesting that the outlier regions were targets of recurrent selection throughout the radiation. Under recurrent sweeps, however, D_xy_ levels of outliers are expected to be lower than those of the genomic background (69); yet D_xy_ levels of outlier regions in *Metrosideros* were significantly elevated. Elevated D_xy_ were even seen in comparisons with a population from the oldest island of Kauai, suggesting that these genomic regions were involved in ancient barriers to gene flow in the common ancestor, or that they harbor ancient divergent haplotypes that predate colonization of the islands (68). Long-term balancing selection or diversifying selection could have maintained increased D_xy_ within differentiation outlier regions as taxa radiated across the islands. Functionally, the enrichment of genes relating to immunity response within these differentiated regions is consistent with the hypothesis that archipelago-wide, shared balancing selection is driving the formation of these regions in *Metrosideros* (88).

Differentiated genomic regions that were lineage specific–and thus likely shaped by taxon-specific divergent selection–were also enriched for ancient variation. This finding, coupled with results from the shared differentiation outlier regions, indicates that both historical and contemporary diversification within *Metrosideros* involved selection on old divergent haplotypes. Several adaptive radiations have been shown to be facilitated by genetic variation that is older than the radiations themselves (23, 25). Selection on old variation may be a common evolutionary mechanism for both animals and plants during adaptive radiation, and may explain how, in remote islands with limited gene pools, lineages are able to repeatedly radiate into new species (15). Utilizing old variation may explain how *Metrosideros*, a long-lived tree, has been able to rapidly and repeatedly colonize novel environments, either through selection on standing ancestral variation (28, 89) or through admixture of ancient divergent haplotypes among hybridizing taxa (33).

The maintenance of ancestral polymorphisms in Hawaiian *Metrosideros* may be aided by the occurrence of a single widespread “generalist” taxon coupled with the rapid evolution of volcanic islands. *M. polymorpha* var. *glaberrima* (G), the only *Metrosideros* taxon found on all of the main islands, has high genetic variation within populations (48) and weak genetic differentiation across the islands, likely playing a role in the origin of the many island-endemic species and varieties of *Metrosideros* (35). Observations on volcanically active Hawaii Island suggest that taxon G is the second taxon to colonize each new island, following rapidly behind early-successional *M. polymorpha* var. *incana* (I). Taxon G maintains the largest population, broadest elevation range, and most generalist niche of any Hawaii Island taxon, and as a result overlaps and hybridizes with all other (habitat-specialist) *Metrosideros* taxa on the island (48). Observations on the extinct volcanoes of the older islands reveal that where divergent selection wanes due to the cessation of lava flows, the differentiation between G and I that is observable on Hawaii Island disappears (35). More broadly, this results suggests that increased introgression would be expected between any co-occurring taxa that are isolated by incomplete intrinsic barriers (52) whenever divergent selection weakens as volcanoes subside and erode. Retention of ancient polymorphisms within taxon G through its massive population size and hybridization with habitat specialists is consistent with the transporter process model proposed by Schluter and Conte (90) and expanded to whole radiations by Martin and Richards (15). Alternating periods of divergent selection across heterogeneous environments and introgression where selection wanes on rapidly evolving volcanic islands would promote linkage among adaptive alleles. In the complex landscape of volcanic islands, this process could lead to the maintenance of linkage disequilibrium among alleles conferring adaptation to a range of environments (i.e., axes of ecological diversification), thus facilitating continued rapid radiation on each new island (15). Future research dissecting the genetic basis of the adaptive phenotype (*e.g*. GWAS (91) or QTL approaches (86)) could clarify the functionality of the ancient variations discovered in the current study and test the importance of the transporter process in the Hawaiian *Metrosideros* radiation.

## Materials and Methods

Detailed description of materials and methods can be found in SI Appendix. Nanopore sequencing data are available from NCBI bioproject ID PRJNA670777. The population genomic sequencing data are available from NCBI bioproject ID PRJNA534153, specifically with the SRR identifiers SRR12673403 to SRR12673495. Data generated from this study, including the reference genome assembly, gene annotation, variant call file, and population genetics statistics, can be found at Zenodo data repository (https://doi.org/10.5281/zenodo.4264399).

## Acknowledgements

We thank the Hawaii Department of Forestry and Wildlife for permission to collect leaf samples from state forests. We also thank Jennifer Johansen, Yohan Pillon, Melissa Johnson, and Chrissen Gemmill for assistance with field collections, Tomoko Sakishima for assistance with greenhouse sample collection and DNA extractions, the College of Agriculture, Forestry, and Natural Resource Management at the University of Hawaii Hilo for greenhouse space, and Angalee Kirby for greenhouse management. We are also grateful to the Genomics Core Facility at Princeton University for sequencing support and the New York University IT High Performance Computing for supplying the computational resources, services, and staff expertise. This research was funded by National Science Foundation Plant Genome Research Program (IOS-1546218), the Zegar Family Foundation (A16-0051), and the NYUAD Research Institute (G1205) to MDP, and the National Science Foundation Faculty Early Career Development Program (DEB0954274) (PI) and Centers of Research Excellence in Science and Technology Program (HRD-0833211) (co-PI) to EAS.

