## Supplemental Materials for "Selection on old variants drives adaptive radiation of *Metrosideros* across the Hawaiian Islands"

**Supplementary Information Text**

**Plant material for the reference genome.** About 25 g of young leaves were collected from a mature individual of *M. polymorpha* var. *incana* (NG4) maintained in a coldframe at the University of Hawaii Hilo. NG4 was produced through a controlled-cross between two trees of this variety occurring along Kuliouou Trail, Oahu (approximate location: 21.3160, -157.7296). Leaf material was collected in small batches, wrapped in aluminum foil, and submerged in liquid $N_2$ within one minute of collection. The bundled samples were then kept at -80C for 72 hours and shipped overnight on dry ice to Oxford Nanopore Technologies, New York, NY and stored again at -80C. The entire tree was covered in a black plastic bag (dark-treated) for 24 hours prior to collection.

**Nanopore sequencing-based whole-genome-, RNA-, and Pore-C sequencing.** Using the Qiagen DNeasy Plant Mini Kit, DNA was extracted from 2 g of collected leaf tissue. Separately, total RNA was extracted from 1 g of collected leaf tissue using Thermofisher's PureLink RNA Mini Kit. Full-length cDNA was synthesized from 50 ng of total RNA using the Oxford Nanopore Technologies PCS109 kit, followed by 14 rounds of PCR amplification using the primer mixture from the Oxford Nanopore Technologies EXP-PCA001 kit. A detailed step-by-step protocol outlining the Pore-C method can be found in our online data repository at https://doi.org/10.5281/zenodo.4264399.

A sequencing library was prepared using the Oxford Nanopore Technologies standard ligation sequencing kit SQK-LSK109. Sequencing was conducted on a GridION X5 sequencer for 72 hours, and the raw data were base-called by Oxford Nanopore Technologies basecaller Guppy (available on https://community.nanoporetech.com/) ver. 3.2.8 for the genomic DNA and Pore-C DNA, and ver. 3.2.10 for the cDNA in the high-accuracy mode.

**Nanopore sequence-based reference genome assembly.** The FASTQ files from the whole-genome sequencing data were filtered for high-quality long reads. We used the program filtlong (https://github.com/rrwick/Filtlong) with parameters --min_length 10000 --min_mean_q 85 --min_window_q 70, which selects for reads longer than 10 kbp, average Q-score greater than 8.2, and a minimum sliding window Q-score of 5.2. The filtered nanopore reads were then assembled with the genome assembler flye (1).

**Assembly contig scaffolding with Pore-C sequencing data.** The Pore-C data analysis was conducted using the Pore-C workflow developed by Oxford Nanopore Technologies (https://github.com/nanoporetech/Pore-C-Snakemake), which uses the snakemake workflow engine (2). Briefly, the workflow first aligns the nanopore Pore-C chromosome contact sequence reads to the unscaffolded *Metrosideros* genome assembly using bwa-sw ver. 0.7.17-r1188 (3) with parameters -b 5 -q 2 -r 1 -T 15 -z 10. Compared to conventional Hi-C data, Pore-C contains an enrichment of higher order contacts (4),and to process the multi-contact nanopore reads we used Pore-C tools (https://github.com/nanoporetech/pore-c) also developed by Oxford Nanopore Technologies. The alignment BAM file was processed with Pore-C tool to filter spurious alignments, detect ligation junctions, and assign fragments that originated from the same chromosomal contacts. Pore-C tools converted the alignment BAM file to a chromosome contact pairs format (https://github.com/4dn-dcic/pairix/blob/master/pairs_format_specification.md) for compatibility with the conventional downstream chromosome contact based analysis methods. The pairs file was converted to a hic file format using the Juicer ver. 1.14.08 tools (5) to use as the input data for the Juicebox assembly tools (6). The Juicebox tools were used to scaffold the draft *Metrosideros* assembly, and we followed established guidelines (https://github.com/theaidenlab/Genome-Assembly-Cookbook) to manually construct the chromosome-scale scaffolds using the Pore-C based chromosome contact frequency information. We assigned chromosome numbers to the superscaffold through synteny with the *Eucalyptus grandis* genome assembly (7). Synteny between the *Metrosideros* superscaffold and *Eucalyptus* chromosomes was determined by aligning the assemblies to each other and visualizing the alignment through the program D-GENIES (8).

**Genome annotation.** The cDNA library that was sequenced on the nanopore sequencer was used to annotate the coding sequence regions for the *M. polymorpha* genome assembly. Initially, we used Pychopper ver. 2.3.1 (https://github.com/nanoporetech/pychopper) to trim primers, identify full-length cDNA sequences, and orient the sequence to the correct strand. A total of 12,298,201 (70.2%) reads were classified by Pychopper and were used for downstream analysis.

The long reads were aligned to the reference genome using minimap2 ver. 2.17-r941 (9) with options -ax splice -uf -k14. The alignment file was then used by stringtie2 ver. 2.1.3b (10), which is optimized for *de novo* transcriptome assembly using long-read sequencing and a reference genome. We used the MAKER program (11) for gene annotation using the workflow outlined on the website https://gist.github.com/darencard/bb1001ac1532dd4225b030cf0cd61ce2. The transcriptome assembly from stringtie2 was used as EST evidence in MAKER, and the protein sequences from the previous *M. polymorpha* assembly (12), *E. grandis* assembly (7), and *A. thaliana* (TAIR10) were used for a protein homology search in MAKER. After an initial round of MAKER annotation the gene models were used by SNAP (13) and Augustus (14) to create gene model training datasets specifically for our *M. polymorpha* genome assembly. The training dataset was used for a second round of MAKER gene annotation.

We identified the repetitive regions of the *M. polymorpha* reference genome, first using Repeatmodeler ver. 1.0.10 (http://www.repeatmasker.org/RepeatModeler/) for the *de novo* identification repeat sequences in the reference genome, and then using Repeatmasker ver. 4.1.0 (http://www.repeatmasker.org/RepeatMasker/) to identify the genomic locations of the repetitive sequences in our reference genome.

***Metrosideros* population sequencing.** For the population genomic sampling, we collected young leaf tissue from 9-11 adults from each of eight taxa on the focal island of Oahu and fewer adults of two taxa on Kauai. Collected leaf tissue was kept cool and stored at -80C within 48 hours of collection. Leaf material from the three outgroup samples was silica-dried in the field and stored in a dessicator jar. DNA was extracted from both frozen and dried leaf samples using the Macherey-Nagel NucleoSpin Plant II Mini kit. We used a Tn5 transposase-based method to prepare the whole-genome sequencing library. Mosaic End adaptor A and B (Tn5ME-A: TCGTCGGCAGCGTCAGATGTGTAT AAGAGACAG; Tn5ME-B: GTCTCGTGGGCTCGGAGATGTGTATAAGAGACAG) was annealed with Rev (Tn5ME-Rev: /5Phos/CTGTCTCTTATACACATCT) by mixing 10uL (100uM) of each oligonucleotide with 80 uL of reassociation buffer (10 mM Tris pH 8.0, 50 mM NaCl, 1 mM EDTA) in BioRad thermocycler with the following program: 95°C for 10 min, 90°C for 1 min, and decrease temperature by 1°C/cycle for 60 cycles, held for 1 min at each temperature. Pre-charge of Tn5 with adapters was carried out in solution by mixing 22.5 uL of 100 ng/uL Tn5 (Tn5 protein was produced following the protocol described by (15)), 76.5 uL reassociation buffer/glycerol (1:1), and 4.5 uL of equal molar of annealed adaptor 1 (A-Rev) and annealed adapter 2 (B-Rev). The reaction was then incubated at 37°C for 30 min. The annealed adapters bind to Tn5 transposase to form the transposome complex.

Genomic DNA was tagmented by mixing with 1 uL of the above assembled Tn5 transposome, 4 uL of 5 X TAPS buffer (50 mM TAPS-NaOH pH 8.5 [Alfa aesar # J63268], 25 mM MgCl2, 50% v/dimethylformamide [ThermoFisher #20673], pH 8.5 at 25°C) and water to a total volume of 20 uL and incubated at 55°C for 7 min. The transposome fragments and attaches adapters to gDNA. The reaction was completed by adding 5 uL of 0.2% SDS (Promega, #V6551) to each reaction followed by incubation at 55°C for 7 min to inactivate and release the Tn5. To enrich the DNA fragments that have adapter molecules on both ends we attached an index to the library: 2µl of the stopped tagmentation, 1µl i5 index primer (1uM), 1µl i7 index primer (1µM), 10µl of OneTaq HS Quick-Load 2x master mix (NEB #M0486L), and 6µl of water were combined to make a 20-ul final reaction. The reaction was heated at 68°C for 3min and 95°C for 30 sec, then thermocycled 12 times at 95°C for 10 sec, 55°C for 30 sec, and 68°C for 30 sec, followed by a final extension of 5 min at 68°C.

We then pooled the libraries together, taking 5uL from individual libraries. The pooled library was cleaned and size-selected using Agencourt AMPure XP beads (Beckman Coulter, #A63881) at a 0.8:1 (beads: DNA) ratio. The final library was quantified with a Qubit high-sensitivity DNA kit (Invitrogen Q32854) and examined on an Agilent 2100 Bioanalyzer high-sensitivity DNA chip (Agilent p/n# 2938-85004) to observe the library size distribution. The sequencing library was loaded on a NovaSeq 6000 S1 flow cell and sequenced under a 2×150-bp conformation at the Genomics Core Facility within the Lewis-Sigler Institute for Integrative Genomics at Princeton University.

**Calling genome-wide polymorphisms.** Raw sequencing reads were downloaded from our previous study (16) and combined with the newly generated sequencing data from the current study. From the sequence read archive (SRA) website we downloaded FASTQs with SRR identifiers SRR8943660 to SRR8943653. The sequencing reads were adapter-trimmed and quality-controlled using BBTools (https://jgi.doe.gov/data-and-tools/bbtools/) bbduk program version 37.66 with option: minlen = 25 qtrim = rl trimq = 10 ktrim = r k = 25 mink = 11 hdist = 1 tpe tbo.

Sequencing reads were then aligned to the scaffolded *M. polymorpha* reference genome generated from this study using bwa-mem. PCR duplicate reads were removed using picard version 2.9.0 (http://broadinstitute.github.io/picard/). Genome-wide read coverage statistics were calculated using GATK version 3.8–0 (https://software.broadinstitute.org/gatk/).

We used the GATK HaplotypeCaller engine to call variant sites from the BAM alignment file for each sample. The option –ERC GVCF was used to output the variants in the gVCF format, and the gVCFs of each sample were merged together to allow a multi-sample joint genotype procedure using GATK GenotypeGVCFs engine. Using standard GATK best-practice hard-filter guidelines, the VariantFiltration engine was used to filter out low-quality polymorphisms. In addition, we removed SNPs that were within 5 bp of an INDEL and polymorphic sites that had less than 80% of individuals with a genotype call.

**Population relationship analysis.** The alignment BAM files were used to analyze the population relationships between samples. We used ANGSD version 0.929 (17) and ngsTools (18) to analyze the genotype likelihoods of each sample and infer the population relationships using a probabilistic framework. We only analyzed potential variant sites where more than 80% of the individuals had a genotype, while enforcing a total sequencing coverage filter such that included sites had a minimum of 1/3 the average total sequencing depth (734×) and a maximum of three times the average total sequencing depth (6,613×). To minimize the effect of linkage on inferences on population relationships, polymorphic sites were randomly pruned using a 10-kbp sliding window with a minimum distance of 5 kbp between random sites.

NGSadmix (19) was used to estimate the admixture proportions (K) for each individual. For each of K = 3 to 15, the analysis was repeated 100 times and the run with the highest log-likelihood was chosen. Phylogeny reconstruction was conducted by estimating the pairwise genetic distances between samples using NGSdist (20), and the genetic distances were used by FastME ver. 2.1.5 (21) to build a neighbor-joining tree. Support of the phylogenetic tree was examined by generating 100 bootstrapped datasets using NGSdist.

**Investigating reticulate evolutionary history.** The genotype call dataset was used to examine the reticulate evolutionary history of *Metrosideros*. We conducted the ABBA-BABA D test (22, 23) and $f_3$ test (24) using the R package admixr (25), which is based on the ADMIXTOOLS suite (26). Variants were polarized using the high-coverage Fiji sample *M. vitiensis* as the outgroup genome.

**Demographic modeling.** We used the methods δaδi (27), G-PhoCS (28), and MSMC (29, 30) to infer the demographic history of Hawaiian *Metrosideros*. For all analyses we used the genotype call dataset.

For δaδi analysis, we initially randomly thinned the dataset picking a SNP every 10 kbp using PLINK ver. 2.0 (31). The site frequency spectrum was estimated using the easySFS.py (https://github.com/isaacovercast/easySFS) script while using Fiji *M. vitiensis* as the outgroup genome and polarizing the polymorphic sites. The easySFS.py script was also used to project down the sample size to maximize the number of sites analyzed. The unfolded site frequency spectrum data were used as input for δaδi, and we fit 20 demographic models (Fig. S9). We optimized the model parameter estimates using the Nelder-Mead method by randomly perturbing the parameter values for four rounds. The parameter estimates were perturbed threefold, twofold, twofold, and onefold in incremental rounds. Each round the perturbation was conducted for 10, 20, 30, and 40 replicates. Demography parameters were extracted from the round with the highest likelihood. Demographic models were compared using Akaike Information Criteria (AIC) values. The δaδi analysis scripts were based on the study by (32).

To prepare our dataset for G-PhoCS analysis we first partitioned our reference genome into 1-kbp loci and determined those that are close to neutrality. Neutral loci were determined by selecting loci that were 5 kb away from a genic sequence, 500 bp away from a repetitive DNA sequence, and at least 10 kbp away from each other. Since G-PhoCS is designed to analyze the variation within a single genome, we selected a single individual with high genome coverage to represent each island. Selected samples included: H207 from Hawaii Island (population $G_{H1}$), X83 from Molokai (population $G_M$), O385 from Oahu (taxon M), and K283 from Kauai (population $G_K$). The *M. vitiensis* sample from Fiji was used as the outgroup. We used G-PhoCS ver. 1.2.3 and ran every demographic model five times to check for convergence in the demographic parameter estimates. Each MCMC run had 1,000,000 iterations, and the initial

500,000 iterations were discarded as burn-in. Priors were modeled using a gamma distribution ($\alpha = 1$ and $\beta = 10{,}000$ for population size and divergence time; $\alpha = 0.002$ and $\beta = 0.00001$ for migration rates). Different demographic models involved fitting migration bands between two terminal lineages. After the MCMC run was complete, the program Tracer version 1.6 (http://tree.bio.ed.ac.uk/software/tracer/) was used to estimate the 95% highest posterior density for each demography parameter. The G-PhoCS-estimated divergence time $\tau$ is scaled according to the mutation rate ($\mu$). To convert divergence time to absolute divergence time T (in years) we used the following equation:

$$T = \frac{\tau \times g}{\mu}$$

where g represents the generation time.

MSMC2 was used to estimate the past changes in effective population sizes and the divergence times between individuals. From each taxon/population we chose a single representative individual for the MSMC2 analysis as follows: O310 (taxon B), O65 (taxon C), O72 (taxon I), O194 (taxon L), H271 (taxon N), O464 (taxon R), O145 (taxon T), plus the individuals used in the G-PhoCS analysis. We used the alignment BAM file for each individual and the mpileup function of samtools ver. 1.3.1 (33) to detect sites that had a minimum base score of 30, a mapping quality score of 30, and the coefficient to downgrade mapping qualities for excessive mismatches at 50. The resulting text pileup output was used by bcftools ver. 1.3.1 to call variant sites but excluding INDELs and limiting the calls to biallelic SNPs. The bamCaller.py script that was provided by the MSMC suite (https://github.com/stschiff/msmc-tools) was then used to produce the per-chromosome masks and VCF files for each individual. A genome-wide mask file was also created for each individual by using the SNPable workflow (http://lh3lh3.users.sourceforge.net/snpable.shtml) on each superscaffold/chromosome assembly and then converting it to BED format using the makeMappabilityMask.py script within the MSMC suite. Phasing was done by using the output from the beagle analysis. The input files for MSMC2 were generated using the generate_multihetsep.py script, which is also part of the MSMC suite. To estimate changes in effective population size we examined the two haplotypes of each individual, while cross coalescence rates were estimated from four haplotypes from two individuals. The combineCrossCoal.py script from the MSMC suite was used to produce the outputs for plotting.

**Population genomic analysis.** We used the gVCFs that were called from the previous step to create a VCF file that had genotype calls for all sites including the non-variant positions. The GATK GenotypeGVCFs engine was used with the option –includeNonVariantSites. We analyzed a population VCF that had both variant and non-variant sites in order to obtain the correct number of sites to be used as the denominator for the population genetic statistics we were calculating. We used the genomics_general package to calculate $\theta$, $D_{xy}$, and $F_{ST}$ in 10 kbp windows, sliding the window by 5 kbp. For each window we imposed a quality filter only analyzing sites that had a minimum quality score of 30 and a minimum depth of 5×. Windows that had more than 30% of the sites with a genotype call after the filter were chosen for downstream analysis.

To search for genomic outliers of differentiation the $F_{ST}$ values were *Z*-transformed ($zF_{ST}$), and genomic windows with $zF_{ST} \geq 4$ were considered outliers (34). To search for genomic regions with lineage-specific accentuated differentiation we used a comparative population genomic approach and to avoid analyzing differentiation outlier regions shaped by linked selection effects (35). For each phylogenetic sister pair, a $zF_{ST}$ genome scan was conducted, and the zFst value at each 10 kbp window was compared with those for each of the three other sister pairs. Genomic windows for which the focal sister pair had $zF_{ST} \geq 4$ but the other sister pairs had $zF_{ST} < 4$ were considered to be regions of lineage-specific accentuated differentiation. Further, we required at least two consecutive windows with evidence of lineage-specific accentuated differentiation to avoid spurious outliers.

The strength of evidence of selective sweeps was estimated using the $\omega$ statistic (36). We used the program OmegaPlus (37), which is specifically designed to estimate the $\omega$ statistics for genome-wide SNP datasets. We set the grid size of OmegaPlus so that the $\omega$ statistics would be estimated at 10-kbp windows for each superscaffold/chromosome.

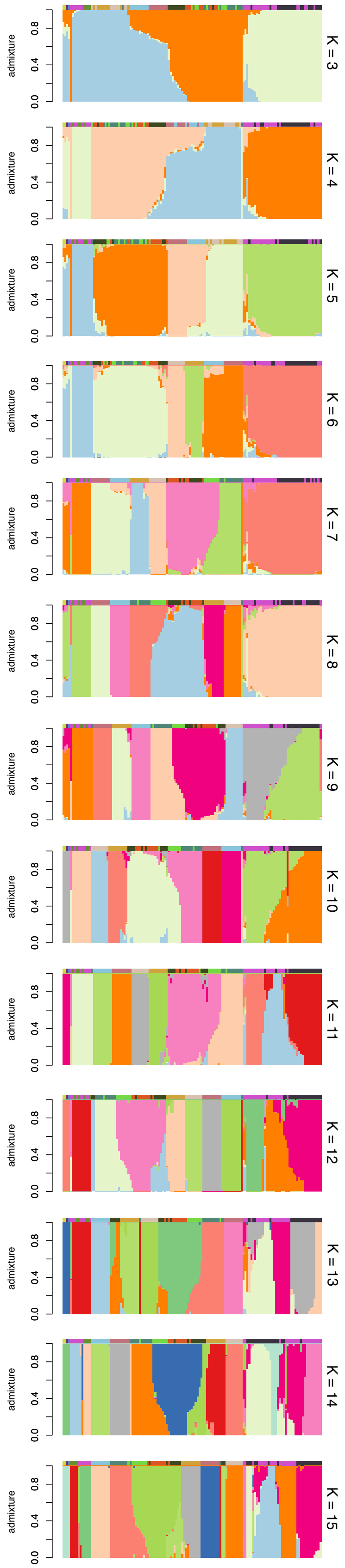


**Fig. S1.** Ancestry proportions for K=3 to K=15 across all Hawaiian archipelago samples.

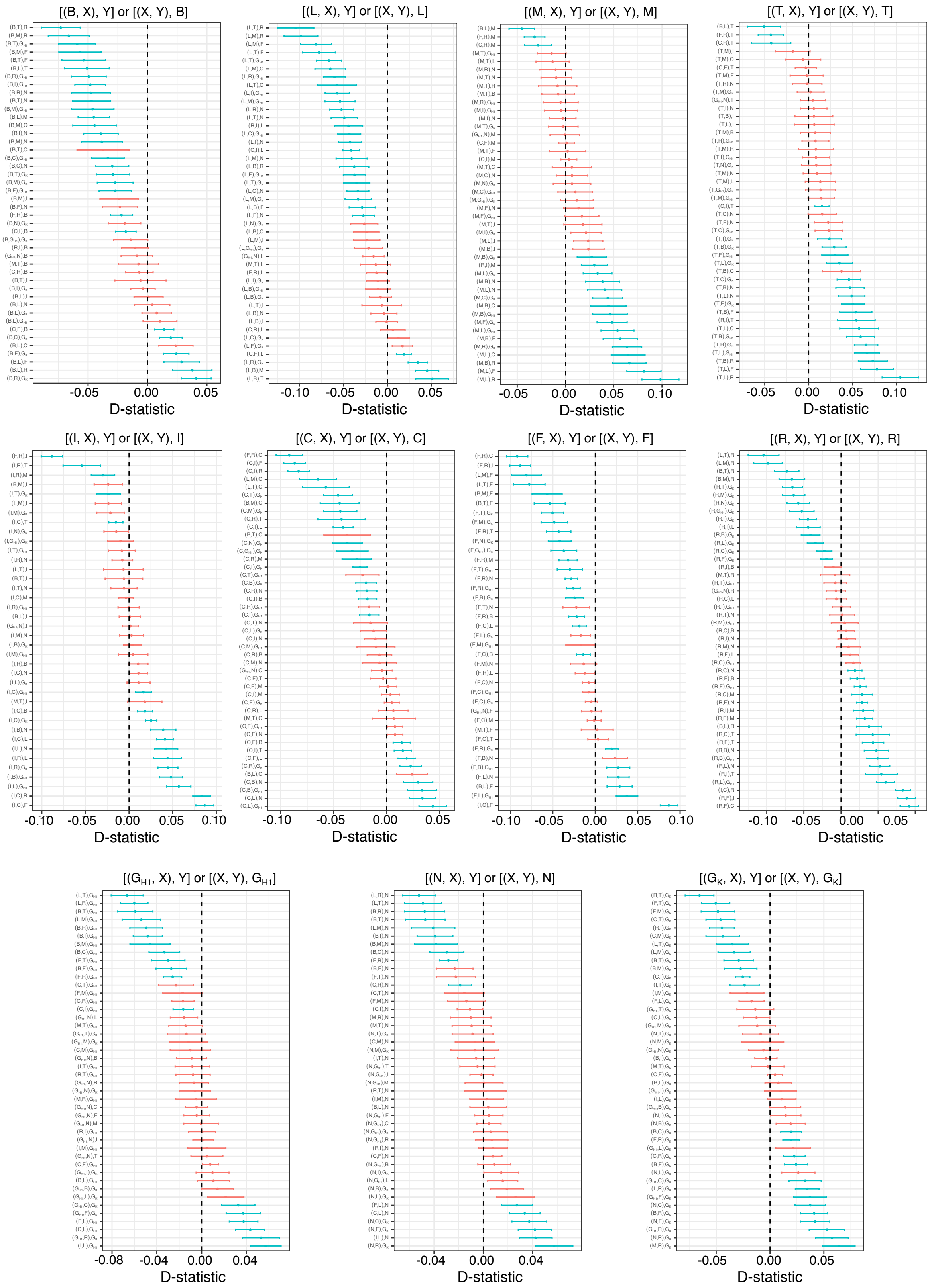


**Fig. S2.** ABBA-BABA D-test statistics for all taxa/population trio combinations.

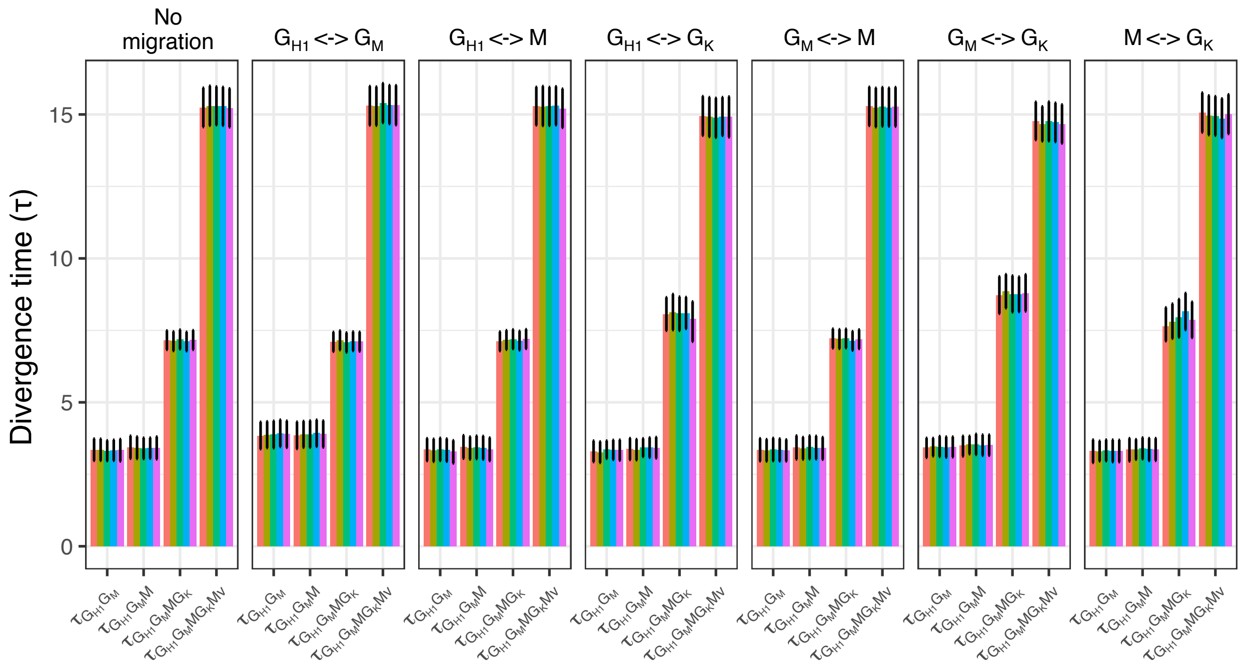


**Fig. S3.** G-PhoCS-based divergence time estimates for different migration models.

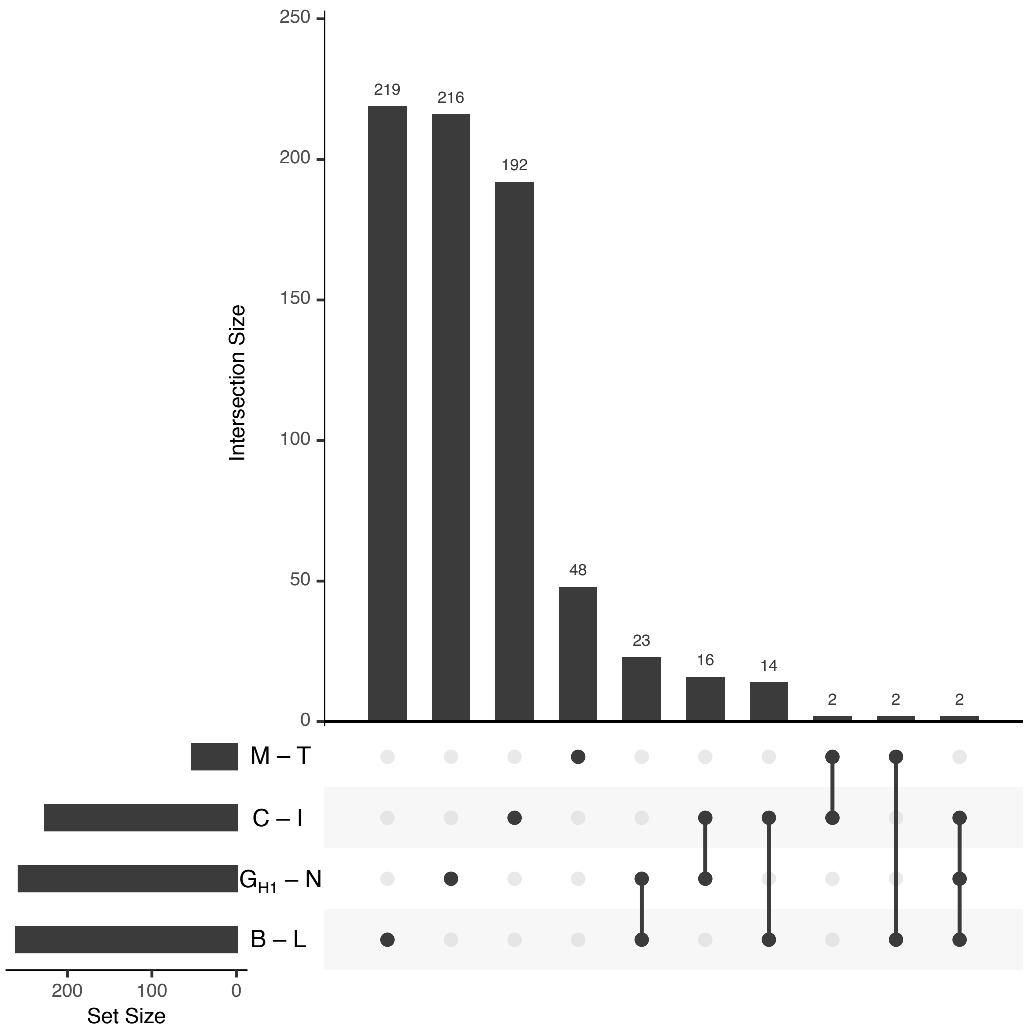


**Fig. S4.** The number of overlapping $zF_{ST}$ outlier positions.

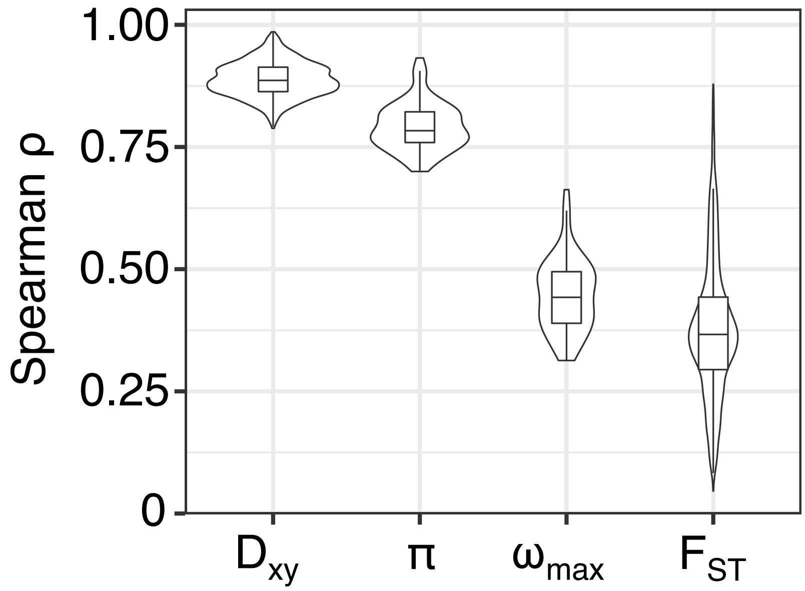


**Fig. S5.** Between-taxon/population correlations in sequence diversity ($\pi$), divergence ($D_{xy}$), differentiation ($F_{ST}$), and selective sweep statistics ($\omega_{max}$).

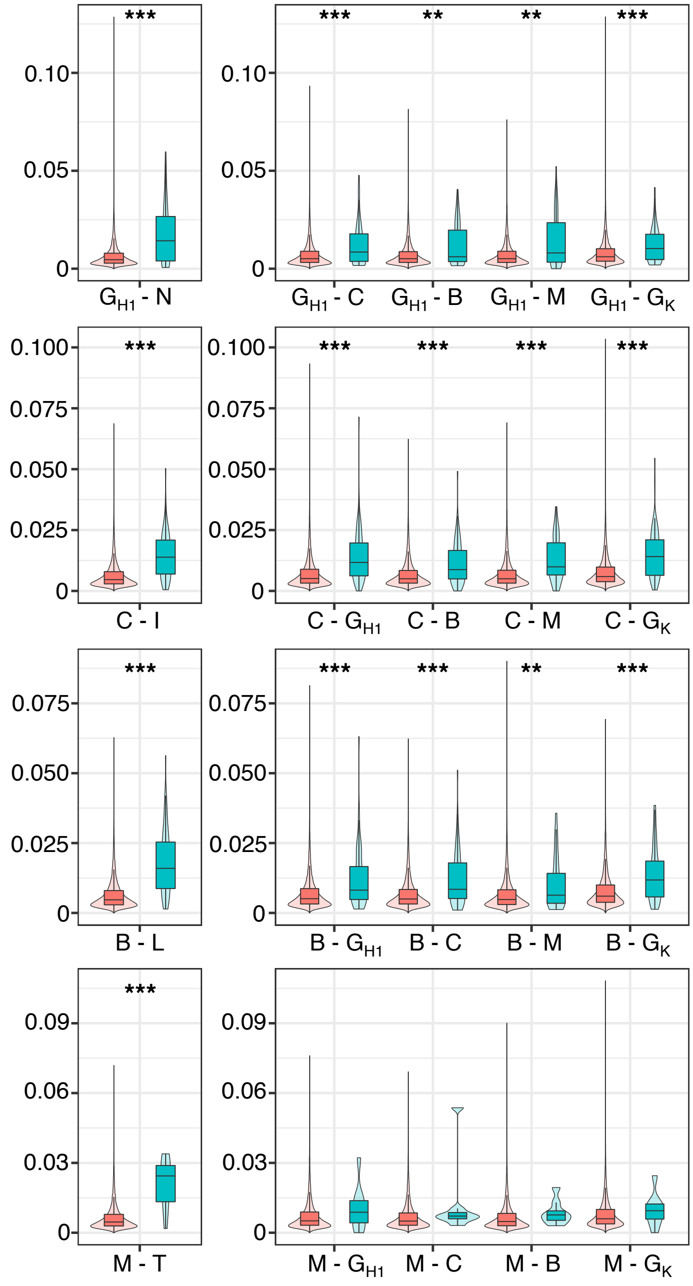


**Fig. S6.** Divergence ($D_{xy}$) statistics identified in a sister pair (left window) and the same pairwise statistics calculated for the same regions between one member of the pair and 4 other taxa representing increasing genomic divergence (right window). Red boxes are statistics from the genomic background, and green boxes are statistics from the Lineage specific accentuated differentiation outlier regions. * indicate $p < 0.05$, ** indicate $p < 0.01$, and *** indicate $p < 0.001$.

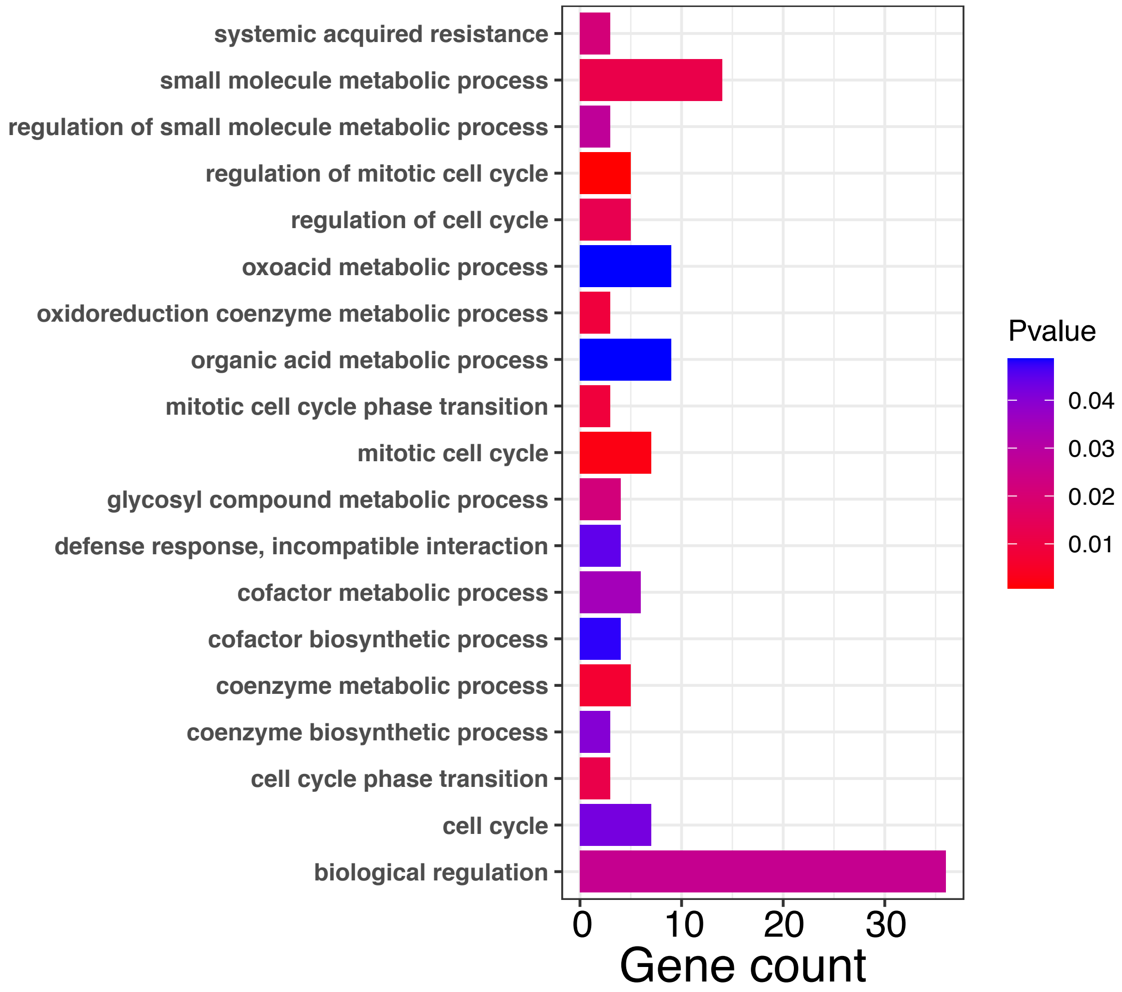


**Fig. S7.** Significantly enriched gene ontology terms for genes overlapping genomic regions of differentiation outliers.

**Table S1.** Sequencing statistics from the nanopore sequencing of the whole genome.

| Flowcell | Number of reads | Median read length | Read length N50 | Median quality score (QS) | Total bases |
|---|---|---|---|---|---|
| | | | Raw | | |
| FAL75935 | 1,821,430 | 2,929 | 13,392 | 11.8 | 11,244,606,349 |
| FAL76650 | 1,909,759 | 2,895 | 13,081 | 11.8 | 11,574,820,609 |
| FAN08536 | 483,618 | 22,785 | 29,641 | 12.0 | 11,013,147,889 |
| Total | 4,214,807 | 3,433 | 18,983 | 11.8 | 33,832,574,847 |
| | | | Filtered | | |
| FAL75935 | 294,849 | 16,568 | 19,761 | 12.5 | 5,458,999,683 |
| FAL76650 | 301,870 | 16,519 | 19,679 | 12.4 | 5,574,321,352 |
| FAN08536 | 282,078 | 25,613 | 29,484 | 12.6 | 7,694,845,247 |
| Total | 878,797 | 19,185 | 23,559 | 12.5 | 18,728,166,282 |

**Table S2.** Aggregated Pore-C sequencing and number of contact results.

| Reads | | | | Read contact order | | | | | | Pairwise Contacts |
|---|---|---|---|---|---|---|---|---|---|---|
| Num of reads | Read bp | Num of reads > 1 contacts | Read bp > 1 contacts | <1 | 2 | 3 | 4 | 5 | 6> | Count |
| 2,562,228 | 5,255,145,653 | 1,523,754 | 3,406,299,103 | 1,038,474 | 615,983 | 426,185 | 232,250 | 119,043 | 130,293 | 7,780,611 |

**Table S3.** Sequencing statistics from the nanopore sequencing of the cDNA library.

| Sequencing statistics | |
|---|---|
| Number of reads | 12,235,945 |
| Total bases | 12,251,510,900 bp |
| Median read length | 847 bp |
| Read length N50 | 1,174 bp |
| Median quality score (QS) | 11.1 |

**Table S4.** Population sample information.

| Sample ID | Code | TaxonName | Island | SourceLocation | Mean Coverage | Median Coverage | Percent Read Aligned | Sequenced By | Latitude | Longitude | Altitude |
|---|---|---|---|---|---|---|---|---|---|---|---|
| K177 | $G_K$ | M. polymorpha var. glaberrima | Kauai | Kuilau Trail, Kaimana, Kauai, HI | 15.29 | 14 | 0.92 | This_study | | | |
| K283 | $G_K$ | M. polymorpha var. glaberrima | Kauai | Iliau Loop Trail, Edge of Waimea Canyon, Kauai, HI | 18.46 | 17 | 0.94 | This_study | | | |
| K287 | $G_K$ | M. polymorpha var. glaberrima | Kauai | Iliau Loop Trail, Edge of Waimea Canyon, Kauai, HI | 15.76 | 14 | 0.95 | This_study | | | |
| K288 | $G_K$ | M. polymorpha var. glaberrima | Kauai | Iliau Loop Trail, Edge of Waimea Canyon, Kauai, HI | 21 | 19 | 0.95 | This_study | | | |
| K290 | $G_K$ | M. polymorpha var. glaberrima | Kauai | Iliau Loop Trail, Edge of Waimea Canyon, Kauai, HI | 13.11 | 12 | 0.94 | This_study | | | |
| K292 | $G_K$ | M. polymorpha var. glaberrima | Kauai | Iliau Loop Trail, Edge of Waimea Canyon, Kauai, HI | 13.35 | 11 | 0.95 | This_study | | | |
| K299 | $G_K$ | M. polymorpha var. glaberrima | Kauai | Iliau Loop Trail, Edge of Waimea Canyon, Kauai, HI | 15.31 | 13 | 0.94 | This_study | | | |
| K202 | WW | M. waialealae var. waialealae | Kauai | Kahili Ridge, Kauai, HI | 7.2 | 5 | 0.91 | This_study | | | |
| K205 | WW | M. waialealae var. waialealae | Kauai | Kahili Ridge, Kauai, HI | 8.21 | 7 | 0.92 | This_study | | | |
| K207 | WW | M. waialealae var. waialealae | Kauai | Kahili Ridge, Kauai, HI | 1.6 | 2 | 0.91 | This_study | | | |
| K214 | WW | M. waialealae var. waialealae | Kauai | Kahili Ridge, Kauai, HI | 14.54 | 12 | 0.96 | This_study | | | |
| O296 | B | M. polymorpha race B | Oahu | Konahuanui, Oahu, HI | 12.42 | 9 | 0.95 | This_study | 21.35 | -157.79 | 953.00 |
| O297 | B | M. polymorpha race B | Oahu | Konahuanui, Oahu, HI | 9.48 | 6 | 0.83 | This_study | 21.35 | -157.79 | 953.00 |
| O301 | B | M. polymorpha race B | Oahu | Konahuanui, Oahu, HI | 16.5 | 14 | 0.92 | This_study | 21.35 | -157.79 | 940.00 |
| O303 | B | M. polymorpha race B | Oahu | Konahuanui, Oahu, HI | 17.07 | 15 | 0.93 | This_study | 21.35 | -157.79 | 929.00 |
| O304 | B | M. polymorpha race B | Oahu | Konahuanui, Oahu, HI | 17.68 | 14 | 0.91 | This_study | 21.35 | -157.79 | 924.00 |
| O305 | B | M. polymorpha race B | Oahu | Konahuanui, Oahu, HI | 16.73 | 14 | 0.95 | This_study | 21.35 | -157.79 | 902.00 |
| O306 | B | M. polymorpha race B | Oahu | Konahuanui, Oahu, HI | 13.35 | 11 | 0.93 | This_study | 21.35 | -157.79 | 896.00 |
| O308 | B | M. polymorpha race B | Oahu | Konahuanui, Oahu, HI | 11.65 | 10 | 0.95 | This_study | 21.35 | -157.79 | 882.00 |
| O309 | B | M. polymorpha race B | Oahu | Konahuanui, Oahu, HI | 16.97 | 13 | 0.93 | This_study | 21.35 | -157.79 | 877.00 |
| O310 | B | M. polymorpha race B | Oahu | Konahuanui, Oahu, HI | 17.79 | 15 | 0.94 | This_study | 21.35 | -157.79 | 871.00 |
| O45 | C | M. polymorpha race C | Oahu | \`Aiea, Oahu, HI | 23.12 | 22 | 0.93 | This_study | 21.42 | -157.86 | 598.00 |
| O55 | C | M. polymorpha race C | Oahu | \`Aiea, Oahu, HI | 20.98 | 20 | 0.90 | This_study | 21.41 | -157.87 | 534.00 |
| O60 | C | M. polymorpha race C | Oahu | \`Aiea, Oahu, HI | 20.81 | 20 | 0.93 | This_study | 21.41 | -157.87 | 499.00 |
| O63 | C | M. polymorpha race C | Oahu | \`Aiea, Oahu, HI | 19.83 | 19 | 0.92 | This_study | 21.41 | -157.87 | 536.00 |
| O64 | C | M. polymorpha race C | Oahu | \`Aiea, Oahu, HI | 31.59 | 30 | 0.92 | This_study | 21.41 | -157.87 | 536.00 |
| O65 | C | M. polymorpha race C | Oahu | \`Aiea, Oahu, HI | 20.22 | 19 | 0.92 | This_study | 21.41 | -157.87 | 537.00 |
| O67 | C | M. polymorpha race C | Oahu | \`Aiea, Oahu, HI | 19.62 | 19 | 0.94 | This_study | 21.41 | -157.87 | 544.00 |
| O68 | C | M. polymorpha race C | Oahu | \`Aiea, Oahu, HI | 18.5 | 18 | 0.93 | This_study | 21.41 | -157.87 | 537.00 |
| O70 | C | M. polymorpha race C | Oahu | \`Aiea, Oahu, HI | 19.06 | 18 | 0.93 | This_study | 21.41 | -157.87 | 527.00 |
| O71 | C | M. polymorpha race C | Oahu | \`Aiea, Oahu, HI | 21.72 | 20 | 0.92 | This_study | 21.41 | -157.87 | 527.00 |
| O119 | F | M. polymorpha race F | Oahu | Konahuanui, Oahu, HI | 9.95 | 9 | 0.90 | This_study | 21.35 | -157.80 | 683.00 |
| O122 | F | M. polymorpha race F | Oahu | Konahuanui, Oahu, HI | 11.04 | 10 | 0.90 | This_study | 21.35 | -157.80 | 677.00 |
| O128 | F | M. polymorpha race F | Oahu | Konahuanui, Oahu, HI | 8.55 | 8 | 0.92 | This_study | 21.35 | -157.80 | 656.00 |
| O131 | F | M. polymorpha race F | Oahu | Konahuanui, Oahu, HI | 14.01 | 11 | 0.89 | This_study | 21.35 | -157.80 | 651.00 |
| O138 | F | M. polymorpha race F | Oahu | Konahuanui, Oahu, HI | 14.8 | 14 | 0.94 | This_study | 21.35 | -157.80 | 658.00 |
| O141 | F | M. polymorpha race F | Oahu | Konahuanui, Oahu, HI | 12.96 | 12 | 0.94 | This_study | 21.35 | -157.80 | 662.00 |
| O142 | F | M. polymorpha race F | Oahu | Konahuanui, Oahu, HI | 13.01 | 12 | 0.92 | This_study | 21.35 | -157.80 | 657.00 |
| O149 | F | M. polymorpha race F | Oahu | Konahuanui, Oahu, HI | 15.14 | 13 | 0.95 | This_study | 21.35 | -157.80 | 648.00 |
| O168 | F | M. polymorpha race F | Oahu | Konahuanui, Oahu, HI | 14.97 | 13 | 0.94 | This_study | 21.35 | -157.80 | 603.00 |
| O72 | I | M. polymorpha var. incana | Oahu | \`Aiea, Oahu, HI | 20.25 | 19 | 0.94 | This_study | 21.41 | -157.87 | 530.00 |
| O75 | I | M. polymorpha var. incana | Oahu | \`Aiea, Oahu, HI | 20.82 | 20 | 0.90 | This_study | 21.41 | -157.88 | 527.00 |
| O76 | I | M. polymorpha var. incana | Oahu | \`Aiea, Oahu, HI | 19.39 | 18 | 0.93 | This_study | 21.41 | -157.88 | 476.00 |
| O77 | I | M. polymorpha var. incana | Oahu | \`Aiea, Oahu, HI | 20.15 | 20 | 0.93 | This_study | 21.41 | -157.88 | 492.00 |
| O78 | I | M. polymorpha var. incana | Oahu | \`Aiea, Oahu, HI | 21.55 | 20 | 0.93 | This_study | 21.41 | -157.87 | 498.00 |
| O88 | I | M. polymorpha var. incana | Oahu | \`Aiea, Oahu, HI | 20.72 | 17 | 0.92 | This_study | 21.41 | -157.87 | 533.00 |
| O89 | I | M. polymorpha var. incana | Oahu | \`Aiea, Oahu, HI | 24.98 | 22 | 0.93 | This_study | 21.41 | -157.87 | 531.00 |
| O90 | I | M. polymorpha var. incana | Oahu | \`Aiea, Oahu, HI | 17.75 | 16 | 0.90 | This_study | 21.41 | -157.87 | 521.00 |
| O91 | I | M. polymorpha var. incana | Oahu | \`Aiea, Oahu, HI | 11.48 | 10 | 0.85 | This_study | 21.41 | -157.87 | 517.00 |
| O92 | I | M. polymorpha var. incana | Oahu | \`Aiea, Oahu, HI | 19.14 | 16 | 0.91 | This_study | 21.41 | -157.87 | 517.00 |
| O177 | L | M. polymorpha race L | Oahu | Mt. Ka\`ala, Oahu, HI | 21.62 | 18 | 0.92 | This_study | 21.51 | -158.14 | 1230.00 |
| O180 | L | M. polymorpha race L | Oahu | Mt. Ka\`ala, Oahu, HI | 13.63 | 11 | 0.93 | This_study | 21.51 | -158.15 | 1224.00 |
| O183 | L | M. polymorpha race L | Oahu | Mt. Ka\`ala, Oahu, HI | 18.4 | 16 | 0.94 | This_study | 21.50 | -158.15 | 1224.00 |
| O189 | L | M. polymorpha race L | Oahu | Mt. Ka\`ala, Oahu, HI | 13.96 | 11 | 0.93 | This_study | 21.50 | -158.15 | 1214.00 |
| O191 | L | M. polymorpha race L | Oahu | Mt. Ka\`ala, Oahu, HI | 16.34 | 13 | 0.95 | This_study | 21.50 | -158.15 | 1211.00 |
| O194 | L | M. polymorpha race L | Oahu | Mt. Ka\`ala, Oahu, HI | 19.09 | 17 | 0.92 | This_study | 21.50 | -158.15 | 1200.00 |
| O196 | L | M. polymorpha race L | Oahu | Mt. Ka\`ala, Oahu, HI | 17.15 | 15 | 0.93 | This_study | 21.50 | -158.15 | 1191.00 |
| O199 | L | M. polymorpha race L | Oahu | Mt. Ka\`ala, Oahu, HI | 14.17 | 12 | 0.95 | This_study | 21.50 | -158.15 | 1202.00 |
| O200 | L | M. polymorpha race L | Oahu | Mt. Ka\`ala, Oahu, HI | 15.44 | 11 | 0.91 | This_study | 21.50 | -158.15 | 1197.00 |
| O355 | L | M. polymorpha race L | Oahu | Mt. Ka\`ala, Oahu, HI | 12.74 | 11 | 0.86 | This_study | 21.50 | -158.15 | 1203.00 |
| NG154 | M | M. macropus | Oahu | Lanihuli, Oahu, HI | 14.45 | 12 | 0.95 | This_study | | | |
| O135 | M | M. macropus | Oahu | Konahuanui, Oahu, HI | 12.65 | 10 | 0.96 | This_study | 21.35 | -157.80 | 658.00 |
| O344 | M | M. macropus | Oahu | Konahuanui, Oahu, HI | 14.57 | 12 | 0.95 | This_study | | | |
| O367 | M | M. macropus | Oahu | Konahuanui, Oahu, HI | 16.59 | 14 | 0.87 | This_study | | | |
| O368 | M | M. macropus | Oahu | Konahuanui, Oahu, HI | 12.75 | 9 | 0.96 | This_study | 21.35 | -157.80 | 579.00 |
| O369 | M | M. macropus | Oahu | Konahuanui, Oahu, HI | 17.3 | 13 | 0.95 | This_study | 21.35 | -157.80 | 659.00 |
| O370 | M | M. macropus | Oahu | Konahuanui, Oahu, HI | 13.03 | 11 | 0.95 | This_study | 21.35 | -157.80 | 659.00 |
| O371 | M | M. macropus | Oahu | Konahuanui, Oahu, HI | 8.85 | 7 | 0.96 | This_study | 21.35 | -157.80 | 657.00 |
| O373 | M | M. macropus | Oahu | Konahuanui, Oahu, HI | 8.43 | 6 | 0.96 | This_study | 21.35 | -157.80 | 663.00 |
| O385 | M | M. macropus | Oahu | Konahuanui, Oahu, HI | 17.14 | 14 | 0.94 | This_study | 21.35 | -157.80 | 657.00 |
| O462 | R | M. rugosa | Oahu | Waikane, Oahu, HI | 10.12 | 9 | 0.89 | This_study | | | |
| O463 | R | M. rugosa | Oahu | Waikane, Oahu, HI | 11.9 | 10 | 0.86 | This_study | | | |
| O464 | R | M. rugosa | Oahu | Waikane, Oahu, HI | 17.81 | 15 | 0.86 | This_study | | | |
| O465 | R | M. rugosa | Oahu | Waikane, Oahu, HI | 10.52 | 9 | 0.77 | This_study | | | |
| O466 | R | M. rugosa | Oahu | Waikane, Oahu, HI | 13.13 | 12 | 0.86 | This_study | | | |
| O467 | R | M. rugosa | Oahu | Waikane, Oahu, HI | 15.48 | 13 | 0.87 | This_study | | | |
| O468 | R | M. rugosa | Oahu | Waikane, Oahu, HI | 14.57 | 12 | 0.84 | This_study | | | |
| O469 | R | M. rugosa | Oahu | Waikane, Oahu, HI | 26.93 | 24 | 0.91 | This_study | | | |
| O470 | R | M. rugosa | Oahu | Waikane, Oahu, HI | 10.22 | 9 | 0.85 | This_study | | | |
| O471 | R | M. rugosa | Oahu | Waikane, Oahu, HI | 9.52 | 8 | 0.82 | This_study | | | |
| M. rugosa | R | M. rugosa | Oahu | | 24.75 | 22 | 0.95 | Choi_etal | - | - | |
| M. tremuloic | T | M. tremuloides | Oahu | | 19.79 | 18 | 0.91 | Choi_etal | - | - | |
| O143 | T | M. tremuloides | Oahu | Konahuanui, Oahu, HI | 12.89 | 12 | 0.91 | This_study | 21.35 | -157.80 | 660.00 |
| O145 | T | M. tremuloides | Oahu | Konahuanui, Oahu, HI | 19.17 | 17 | 0.95 | This_study | 21.35 | -157.80 | 652.00 |
| O146 | T | M. tremuloides | Oahu | Konahuanui, Oahu, HI | 16.62 | 14 | 0.95 | This_study | 21.35 | -157.80 | 652.00 |
| O148 | T | M. tremuloides | Oahu | Konahuanui, Oahu, HI | 14.31 | 13 | 0.93 | This_study | 21.35 | -157.80 | 650.00 |
| O151 | T | M. tremuloides | Oahu | Konahuanui, Oahu, HI | 13.75 | 12 | 0.94 | This_study | 21.35 | -157.80 | 650.00 |
| O154 | T | M. tremuloides | Oahu | Konahuanui, Oahu, HI | 2.45 | 3 | 0.94 | This_study | | | |
| O157 | T | M. tremuloides | Oahu | Konahuanui, Oahu, HI | 18.1 | 16 | 0.94 | This_study | 21.35 | -157.80 | 635.00 |
| O166 | T | M. tremuloides | Oahu | Konahuanui, Oahu, HI | 14.81 | 11 | 0.94 | This_study | 21.35 | -157.80 | 604.00 |
| X36 | $G_M$ | M. polymorpha var. glaberrima | Molokai | | 24.63 | 23 | 0.95 | Choi_etal | 21.14 | -156.93 | |
| X83 | $G_M$ | M. polymorpha var. glaberrima | Molokai | | 13 | 12 | 0.95 | Choi_etal | 21.12 | -156.92 | |
| NG207 | - | M. polymorpha var. glaberrima | Hawaii | Saddle Rd, Hawaii Island, HI | 3.64 | 3 | 0.92 | This_study | | | |
| E106 | N | M. polymorpha var. newellii | Hawaii | | 20.38 | 19 | 0.93 | Choi_etal | 19.72 | -155.19 | |
| E110 | N | M. polymorpha var. newellii | Hawaii | | 18.85 | 18 | 0.95 | Choi_etal | 19.72 | -155.19 | |
| E111 | $G_{H2}$ | M. polymorpha var. glaberrima | Hawaii | | 21.67 | 19 | 0.96 | Choi_etal | 19.72 | -155.19 | |
| E114 | $G_{H2}$ | M. polymorpha var. glaberrima | Hawaii | | 5.89 | 5 | 0.96 | Choi_etal | 19.72 | -155.19 | |
| E126 | $G_{H2}$ | M. polymorpha var. glaberrima | Hawaii | | 27.09 | 23 | 0.97 | Choi_etal | 19.72 | -155.19 | |
| E132 | $G_{H2}$ | M. polymorpha var. glaberrima | Hawaii | | 14.22 | 11 | 0.94 | Choi_etal | 19.71 | -155.19 | |
| E70 | - | M. polymorpha var. glaberrima | Hawaii | | 18.96 | 15 | 0.95 | Choi_etal | 19.71 | -155.24 | |
| E72 | - | M. polymorpha var. glaberrima | Hawaii | | 17.83 | 15 | 0.96 | Choi_etal | 19.71 | -155.24 | |
| E74 | - | M. polymorpha var. glaberrima | Hawaii | | 35.43 | 36 | 0.95 | Choi_etal | 19.71 | -155.24 | |
| H193 | $G_{H1}$ | M. polymorpha var. glaberrima | Hawaii | | 25.85 | 24 | 0.91 | Choi_etal | 20.07 | -155.67 | |
| H195 | - | M. polymorpha var. glaberrima | Hawaii | | 17.32 | 15 | 0.90 | Choi_etal | 20.07 | -155.67 | |
| H196 | - | M. polymorpha var. glaberrima | Hawaii | | 0.04 | 1 | 0.96 | Choi_etal | 20.07 | -155.67 | |
| H197 | $G_{H1}$ | M. polymorpha var. glaberrima | Hawaii | | 46.72 | 43 | 0.96 | Choi_etal | 20.07 | -155.67 | |
| H198 | $G_{H1}$ | M. polymorpha var. glaberrima | Hawaii | | 22.92 | 21 | 0.93 | Choi_etal | 20.07 | -155.67 | |
| H202 | $G_{H1}$ | M. polymorpha var. glaberrima | Hawaii | | 22.7 | 21 | 0.94 | Choi_etal | 20.07 | -155.67 | |
| H204 | $G_{H1}$ | M. polymorpha var. glaberrima | Hawaii | | 10.87 | 9 | 0.95 | Choi_etal | 20.07 | -155.67 | |
| H206 | $G_{H1}$ | M. polymorpha var. glaberrima | Hawaii | | 11.25 | 9 | 0.92 | Choi_etal | 20.07 | -155.67 | |
| H207 | $G_{H1}$ | M. polymorpha var. glaberrima | Hawaii | | 18.87 | 16 | 0.96 | Choi_etal | 20.07 | -155.67 | |
| H212 | $G_{H1}$ | M. polymorpha var. glaberrima | Hawaii | | 15.26 | 14 | 0.93 | Choi_etal | 20.07 | -155.67 | |
| H235 | - | M. polymorpha var. newellii | Hawaii | | 23.2 | 21 | 0.96 | Choi_etal | 19.68 | -155.15 | |
| H261 | - | M. polymorpha var. newellii | Hawaii | | 8.82 | 7 | 0.96 | Choi_etal | 19.71 | -155.23 | |
| H269 | N | M. polymorpha var. newellii | Hawaii | | 20.08 | 19 | 0.96 | Choi_etal | 19.71 | -155.23 | |
| H270 | N | M. polymorpha var. newellii | Hawaii | | 21.01 | 20 | 0.96 | Choi_etal | 19.71 | -155.23 | |
| H271 | N | M. polymorpha var. newellii | Hawaii | | 17.16 | 16 | 0.95 | Choi_etal | 19.71 | -155.23 | |
| H276 | N | M. polymorpha var. newellii | Hawaii | | 23.37 | 22 | 0.96 | Choi_etal | 19.71 | -155.23 | |
| H300 | N | M. polymorpha var. newellii | Hawaii | | 32.62 | 32 | 0.96 | Choi_etal | 19.71 | -155.22 | |
| H314 | N | M. polymorpha var. newellii | Hawaii | | 16.51 | 15 | 0.93 | Choi_etal | 19.71 | -155.21 | |
| H323 | - | M. polymorpha var. newellii | Hawaii | | 21.64 | 18 | 0.96 | Choi_etal | 19.70 | -155.20 | |
| H329 | N | M. polymorpha var. newellii | Hawaii | | 7.18 | 5 | 0.97 | Choi_etal | 19.71 | -155.19 | |
| H355 | N | M. polymorpha var. newellii | Hawaii | | 27.5 | 24 | 0.97 | Choi_etal | 19.71 | -155.15 | |
| H360 | N | M. polymorpha var. newellii | Hawaii | | 27.15 | 24 | 0.97 | Choi_etal | 19.71 | -155.15 | |
| H404 | - | M. polymorpha var. glaberrima | Hawaii | | 15.51 | 13 | 0.95 | Choi_etal | 20.14 | -155.75 | |
| H439 | - | M. polymorpha var. newellii | Hawaii | | 17.97 | 15 | 0.96 | Choi_etal | 20.04 | -155.69 | |
| H469 | N | M. polymorpha var. newellii | Hawaii | | 23.15 | 18 | 0.96 | Choi_etal | 19.89 | -155.17 | |
| H470 | N | M. polymorpha var. newellii | Hawaii | | 28.4 | 24 | 0.97 | Choi_etal | 19.89 | -155.17 | |
| H493 | - | M. polymorpha var. newellii | Hawaii | | 22.96 | 17 | 0.90 | Choi_etal | 20.02 | -155.45 | |
| H504 | - | M. polymorpha var. newellii | Hawaii | | 3.65 | 2 | 0.69 | Choi_etal | 19.85 | -155.16 | |
| H508 | N | M. polymorpha var. newellii | Hawaii | | 24.6 | 15 | 0.96 | Choi_etal | 19.92 | -155.17 | |
| P31 | Me | M. excelsa | - | New Zealand | 3.95 | 3 | 0.97 | This_study | | | |
| P30 | Mr | M. robusta | - | New Zealand | 6.18 | 4 | 0.96 | This_study | | | |
| AS09 | Mv | M. vitiensis | - | American Samoa | 9 | 5 | 0.83 | This_study | | | |
| P22 | Mv | M. vitiensis | - | Fiji | 10.74 | 8 | 0.94 | This_study | | | |

**Table S5.** δaδi results for all 20 demography models for all 4 sister pairs.

| Model | log-likelihoo | AIC | theta | nu1 | nu2 | nu1a | nu2a | nu1b | nu2b | m12 | m21 | m12a | m21a | m12b | m21b | T1 | T2 | T3 |
|---|---|---|---|---|---|---|---|---|---|---|---|---|---|---|---|---|---|---|
| | | | | | | | | | B vs L | | | | | | | | | |
| (K) secondary contact with symmetric migration and population size change | -1322.58 | 2659.16 | 3451.83 | - | - | 0.7786 | 0.5566 | 1.9384 | 1.9225 | 0.514 | - | - | - | - | - | 0.2281 | 0.2918 | - |
| (O) two epoch of symmetric migration | -1357.08 | 2726.16 | 3121.36 | 1.9189 | 1.3714 | - | - | - | - | - | - | 0.0759 | - | 0.7318 | - | 0.4732 | 0.1646 | - |
| (H) secondary contact with asymmetric migration | -1376.12 | 2764.24 | 2947.57 | 2.0169 | 1.5855 | - | - | - | - | 0.4709 | 0.5356 | - | - | - | - | 0.3928 | 0.2939 | - |
| (E) symmetric migration with population size change | -1385.23 | 2784.46 | 3325.77 | - | - | 1.4578 | 0.5669 | 1.9474 | 1.3091 | 0.5088 | - | - | - | - | - | 0.2638 | 0.354 | - |
| (B) symmetric migration | -1417.07 | 2842.14 | 2959.52 | 2.0664 | 1.4611 | - | - | - | - | 0.4322 | - | - | - | - | - | 0.7063 | - | - |
| (G) secondary contact with symmetric migration | -1418.45 | 2846.9 | 2909.33 | 1.9254 | 1.3774 | - | - | - | - | 0.5568 | - | - | - | - | - | 0.2994 | 0.534 | - |
| (R) secondary contact with asymmetric migration under three epoch (isolation - migration - isolation) | -1419.6 | 2853.2 | 2784.57 | 2.043 | 1.7189 | - | - | - | - | 2.3693 | 1.1876 | - | - | - | - | 0.5645 | 0.1166 | 3.5671 |
| (F) asymmetric migration and population size change | -1422.79 | 2861.58 | 2451.21 | - | - | 1.2181 | 0.2806 | 2.2552 | 2.3538 | 0.5323 | 0.2851 | - | - | - | - | 0.1309 | 0.9174 | - |
| (S) secondary contact with symmetric migration population size change under three epoch (isolation - migration - isolation) | -1435.59 | 2887.18 | 2483.93 | - | - | 19.4443 | 0.3535 | 2.1869 | 1.7371 | 0.4786 | - | - | - | - | - | 0.123 | 1.0174 | 0.0116 |
| (C) asymmetric migration | -1440.41 | 2890.82 | 2811.63 | 2.1431 | 1.6868 | - | - | - | - | 0.3678 | 0.3945 | - | - | - | - | 0.7474 | - | - |
| (T) secondary contact with asymmetric migration population size change under three epoch (isolation - migration - isolation) | -1601.04 | 3220.08 | 3212.22 | - | - | 1.8862 | 0.3564 | 2.9301 | 1.6835 | 0.2513 | 3.147 | - | - | - | - | 0.1902 | 0.146 | 0.145 |
| (I) ancestral symmetric migration | -1665.25 | 3340.5 | 2632.89 | 2.0042 | 1.6651 | - | - | - | - | 0.9616 | - | - | - | - | - | 0.879 | 0.131 | - |
| (P) two epoch of asymmetric migration | -1668.47 | 3352.94 | 1177.11 | 4.3647 | 3.0194 | - | - | - | - | - | - | 5.5715 | 0.464 | 0.229 | 0.2377 | 4.2787 | 1.869 | - |
| (J) ancestral asymmetric migration | -1683.79 | 3379.58 | 2973 | 1.6292 | 1.2504 | - | - | - | - | 0.9233 | 1.049 | - | - | - | - | 0.9208 | 0.0639 | - |
| (N) ancestral asymmetric migration with population size change | -1863.83 | 3743.66 | 1653.34 | - | - | 1.8043 | 1.3067 | 5.0066 | 3.2053 | 1.5766 | 3.656 | - | - | - | - | 3.0309 | 0.5117 | - |
| (M) ancestral symmetric migration with population size change | -1922.59 | 3859.18 | 4007.25 | - | - | 0.6065 | 0.4997 | 4.8627 | 3.1101 | 0.3963 | - | - | - | - | - | 0.1158 | 0.131 | - |
| (D) no migration with population size change | -1935.76 | 3883.52 | 3959.14 | - | - | 0.6841 | 0.5264 | 2.7482 | 2.3279 | - | - | - | - | - | - | 0.088 | 0.1584 | - |
| (Q) secondary contact with symmetric migration under three epoch (isolation - migration - isolation) | -1936.34 | 3884.68 | 2730.04 | 1.9826 | 1.1116 | - | - | - | - | 4.4816 | - | - | - | - | - | 0.9534 | 0.1019 | 0.1821 |
| (A) no migration | -1995.35 | 3996.7 | 3715.68 | 1.9358 | 1.3804 | - | - | - | - | - | - | - | - | - | - | 0.3143 | - | - |
| (L) secondary contact with asymmetric migration and population size change | -2270.84 | 4557.68 | 3843.12 | - | - | 0.3238 | 11.7112 | 1.8081 | 0.4678 | 0.2155 | 2.9631 | - | - | - | - | 1.0995 | 0.8536 | - |
| | | | | | | | | | $G_{H1}$ vs N | | | | | | | | | |
| (O) two epoch of symmetric migration | -985.13 | 1982.26 | 3106.12 | 1.5946 | 1.3956 | - | - | - | - | - | - | 0.5827 | - | 0.9817 | - | 0.3535 | 0.2706 | - |
| (R) secondary contact with asymmetric migration under three epoch (isolation - migration - isolation) | -988.35 | 1990.7 | 3145.91 | 2.045 | 1.1846 | - | - | - | - | 2.41 | 7.3335 | - | - | - | - | 0.4663 | 0.0422 | 5.4051 |
| (G) secondary contact with symmetric migration | -1005.59 | 2021.18 | 2911.75 | 1.6754 | 1.5555 | - | - | - | - | 0.848 | - | - | - | - | - | 0.0343 | 0.6724 | - |
| (C) asymmetric migration | -1016.71 | 2043.42 | 2900.22 | 1.78 | 1.5327 | - | - | - | - | 0.6218 | 0.9316 | - | - | - | - | 0.6906 | - | - |
| (K) secondary contact with symmetric migration and population size change | -1052.91 | 2119.82 | 2945.76 | - | - | 1.3458 | 0.3478 | 1.916 | 1.4179 | 0.9676 | - | - | - | - | - | 0.2502 | 0.5832 | - |
| (H) secondary contact with asymmetric migration | -1058.24 | 2128.48 | 3077.18 | 1.2441 | 1.5301 | - | - | - | - | 1.581 | 0.7094 | - | - | - | - | 0.3014 | 0.4243 | - |
| (E) symmetric migration with population size change | -1062.58 | 2139.16 | 1962.7 | - | - | 0.0579 | 3.0444 | 2.4837 | 1.9242 | 0.6876 | - | - | - | - | - | 1.3156 | 1.5752 | - |
| (L) secondary contact with asymmetric migration and population size change | -1120.55 | 2257.1 | 1707.25 | - | - | 0.3682 | 0.2582 | 3.2357 | 1.7831 | 0.379 | 0.9169 | - | - | - | - | 1.7921 | 1.8851 | - |
| (F) asymmetric migration and population size change | -1204.8 | 2425.6 | 7498.76 | - | - | 0.4041 | 0.066 | 0.9611 | 0.4363 | 0.249 | 6.0537 | - | - | - | - | 5.1216 | 0.1719 | - |
| (T) secondary contact with asymmetric migration population size change under three epoch (isolation - migration - isolation) | -1205.72 | 2429.44 | 2093.11 | - | - | 0.0928 | 0.7276 | 2.0026 | 2.2502 | 1.2399 | 0.4926 | - | - | - | - | 0.452 | 1.4495 | 0.034 |
| (M) ancestral symmetric migration with population size change | -1211.01 | 2436.02 | 3211.78 | - | - | 1.216 | 1.7234 | 1.5029 | 1.1368 | 1.488 | - | - | - | - | - | 0.5262 | 0.0634 | - |
| (Q) secondary contact with symmetric migration under three epoch (isolation - migration - isolation) | -1266.08 | 2544.16 | 2205.43 | 2.6816 | 1.6849 | - | - | - | - | 3.8161 | - | - | - | - | - | 1.0274 | 0.128 | 0.1328 |
| (I) ancestral symmetric migration | -1304.92 | 2619.84 | 2999.38 | 1.5383 | 1.4232 | - | - | - | - | 26.1851 | - | - | - | - | - | 0.5518 | 0.1697 | - |
| (P) two epoch of asymmetric migration | -1320.69 | 2657.38 | 2769.47 | 1.2541 | 1.9424 | - | - | - | - | - | - | 5.564 | 0.0645 | 2.1746 | 0.1079 | 0.8375 | 0.3446 | - |
| (J) ancestral asymmetric migration | -1323.57 | 2659.14 | 2318.22 | 2.2324 | 1.4854 | - | - | - | - | 0.3962 | 4.1008 | - | - | - | - | 1.6856 | 0.1306 | - |
| (S) secondary contact with symmetric migration population size change under three epoch (isolation - migration - isolation) | -1351.4 | 2718.8 | 672.5 | - | - | 2.9562 | 0.4088 | 5.4202 | 5.8626 | 0.4472 | - | - | - | - | - | 5.0116 | 5.1573 | 0.251 |
| (B) symmetric migration | -1376.15 | 2760.3 | 1516.92 | 2.9357 | 2.6375 | - | - | - | - | 0.5183 | - | - | - | - | - | 2.651 | - | - |
| (A) no migration | -1526.21 | 3058.42 | 3988.4 | 1.6743 | 1.3762 | - | - | - | - | - | - | - | - | - | - | 0.2008 | - | - |
| (N) ancestral asymmetric migration with population size change | -1529.26 | 3074.52 | 3878.77 | - | - | 7.85 | 26.963 | 1.486 | 1.2973 | 0.4348 | 0.9963 | - | - | - | - | 0.0484 | 0.1748 | - |
| (D) no migration with population size change | -1550.67 | 3113.34 | 3975.28 | - | - | 22.6936 | 1.3213 | 1.0187 | 1.3429 | - | - | - | - | - | - | 0.113 | 0.0963 | - |
| | | | | | | | | | C vs I | | | | | | | | | |
| (O) two epoch of symmetric migration | -1091.03 | 2194.06 | 3024.92 | 0.8638 | 5.8947 | - | - | - | - | - | - | 0.2736 | - | 15.2904 | - | 0.5115 | 0.0592 | - |
| (G) secondary contact with symmetric migration | -1119.13 | 2248.26 | 2915.47 | 1.2173 | 4.5381 | - | - | - | - | 11.4593 | - | - | - | - | - | 0.5176 | 0.0729 | - |
| (E) symmetric migration with population size change | -1156.87 | 2327.74 | 8174.78 | - | - | 0.0859 | 1.8727 | 0.5935 | 1.7333 | 18.0019 | - | - | - | - | - | 12.3888 | 0.1354 | - |
| (P) two epoch of asymmetric migration | -1235.79 | 2487.58 | 2902.19 | 2.3848 | 4.6283 | - | - | - | - | - | - | 0.5042 | 1.004 | 1.4067 | 9.4359 | 0.3944 | 0.2555 | - |
| (C) asymmetric migration | -1253.84 | 2517.68 | 3025.92 | 3.1179 | 3.9395 | - | - | - | - | 0.2878 | 9.4204 | - | - | - | - | 0.5422 | - | - |
| (H) secondary contact with asymmetric migration | -1255.89 | 2523.78 | 3026.62 | 2.3434 | 5.2936 | - | - | - | - | 1.8644 | 8.7638 | - | - | - | - | 0.0121 | 0.5353 | - |
| (B) symmetric migration | -1274.02 | 2556.04 | 3136.09 | 1.4342 | 5.1103 | - | - | - | - | 8.4561 | - | - | - | - | - | 0.4495 | - | - |
| (I) ancestral symmetric migration | -1306.38 | 2622.76 | 3144.11 | 1.6976 | 4.4919 | - | - | - | - | 10.5463 | - | - | - | - | - | 0.4093 | 0.0105 | - |
| (F) asymmetric migration and population size change | -1337.48 | 2690.96 | 1771.62 | - | - | 0.1983 | 1.6391 | 1.9934 | 8.3463 | 7.033 | 1.9232 | - | - | - | - | 6.9754 | 0.7126 | - |
| (J) ancestral asymmetric migration | -1468.33 | 2948.66 | 2638.51 | 1.8252 | 4.7451 | - | - | - | - | 29.6255 | 1.2755 | - | - | - | - | 0.6613 | 0.0232 | - |
| (M) ancestral symmetric migration with population size change | -1506.49 | 3026.98 | 3377.72 | - | - | 1.5991 | 1.0153 | 6.2071 | 24.0289 | 25.5961 | - | - | - | - | - | 0.3761 | 0.0781 | - |
| (K) secondary contact with symmetric migration and population size change | -1537.19 | 3088.38 | 1142.17 | - | - | 0.1781 | 0.1657 | 2.2247 | 23.1174 | 4.0433 | - | - | - | - | - | 2.4777 | 2.8818 | - |
| (S) secondary contact with symmetric migration population size change under three epoch (isolation - migration - isolation) | -1566.83 | 3149.66 | 2134.6 | - | - | 0.6552 | 0.1955 | 1.835 | 7.9273 | 13.5999 | - | - | - | - | - | 0.1319 | 1.0756 | 0.0173 |
| (Q) secondary contact with symmetric migration under three epoch (isolation - migration - isolation) | -1707.18 | 3426.36 | 2301.44 | 1.6228 | 7.5548 | - | - | - | - | 5.367 | - | - | - | - | - | 0.3248 | 0.5493 | 0.0236 |
| (T) secondary contact with asymmetric migration population size change under three epoch (isolation - migration - isolation) | -1716.21 | 3450.42 | 1232.09 | - | - | 5.4939 | 1.1765 | 3.9713 | 7.9346 | 13.9931 | 0.3071 | - | - | - | - | 8.3158 | 2.5292 | 0.0342 |
| (N) ancestral asymmetric migration with population size change | -1876.62 | 3769.24 | 4168.87 | - | - | 3.8477 | 23.0938 | 3.2409 | 6.9858 | 0.5037 | 5.3401 | - | - | - | - | 0.0979 | 0.0287 | - |
| (A) no migration | -1988.48 | 3982.96 | 4224.85 | 4.7308 | 16.0142 | - | - | - | - | - | - | - | - | - | - | 0.1138 | - | - |
| (D) no migration with population size change | -2033.33 | 4078.66 | 4223.78 | - | - | 7.5318 | 14.3295 | 2.3013 | 2.6296 | - | - | - | - | - | - | 0.0968 | 0.0239 | - |
| (R) secondary contact with asymmetric migration under three epoch (isolation - migration - isolation) | -2186.84 | 4387.68 | 3656.55 | 4.7998 | 8.865 | - | - | - | - | 0.5733 | 2.422 | - | - | - | - | 0.1294 | 0.0277 | 0.1471 |
| (L) secondary contact with asymmetric migration and population size change | -2553.1 | 5122.2 | 8182.32 | - | - | 8.2458 | 2.0076 | 0.5881 | 2.4257 | 0.1099 | 22.9295 | - | - | - | - | 20.7734 | 17.8449 | - |
| | | | | | | | | | M vs T | | | | | | | | | |
| (S) secondary contact with symmetric migration population size change under three epoch (isolation - migration - isolation) | -2076 | 4168 | 4199.58 | - | - | 0.3094 | 1.1757 | 2.9057 | 0.9393 | 1.8086 | - | - | - | - | - | 0.3692 | 0.0641 | 0.0619 |
| (M) ancestral symmetric migration with population size change | -2215.89 | 4445.78 | 3822.62 | - | - | 0.5073 | 0.6306 | 15.2277 | 15.4825 | 0.7442 | - | - | - | - | - | 1.3466 | 0.0797 | - |
| (F) asymmetric migration and population size change | -2274 | 4564 | 4225.99 | - | - | 0.272 | 0.8452 | 2.2173 | 0.8006 | 0.917 | 0.4443 | - | - | - | - | 1.615 | 0.1001 | - |
| (H) secondary contact with asymmetric migration | -2449.35 | 4910.7 | 3685.32 | 0.831 | 1.193 | - | - | - | - | 0.7101 | 0.3337 | - | - | - | - | 0.4972 | 0.2582 | - |
| (G) secondary contact with symmetric migration | -2546.86 | 5103.72 | 3635.68 | 0.9581 | 1.1164 | - | - | - | - | 0.538 | - | - | - | - | - | 0.5689 | 0.1793 | - |
| (C) asymmetric migration | -2654.28 | 5318.56 | 4019.14 | 0.7266 | 0.9885 | - | - | - | - | 0.5459 | 0.337 | - | - | - | - | 0.7347 | - | - |
| (B) symmetric migration | -2754.61 | 5517.22 | 3808.84 | 0.8437 | 0.9757 | - | - | - | - | 0.4692 | - | - | - | - | - | 0.8443 | - | - |
| (K) secondary contact with symmetric migration and population size change | -2786.1 | 5586.2 | 3732.87 | - | - | 15.1116 | 1.4335 | 0.8697 | 1.11 | 0.3407 | - | - | - | - | - | 0.0573 | 0.6637 | - |
| (O) two epoch of symmetric migration | -2835.26 | 5682.52 | 2933.45 | 1.0421 | 1.3858 | - | - | - | - | - | - | 0.7857 | - | 0.3321 | - | 0.3907 | 1.0894 | - |
| (P) two epoch of asymmetric migration | -2892.66 | 5801.32 | 1689.53 | 1.609 | 2.5379 | - | - | - | - | - | - | 22.807 | 2.6106 | 0.3055 | 0.1493 | 1.8543 | 2.2994 | - |
| (E) symmetric migration with population size change | -2910.24 | 5834.48 | 2475.92 | - | - | 0.0448 | 0.251 | 1.3998 | 1.7064 | 0.3084 | - | - | - | - | - | 1.3624 | 1.7524 | - |
| (I) ancestral symmetric migration | -3174.54 | 6359.08 | 4076.97 | 0.724 | 0.7996 | - | - | - | - | 0.8232 | - | - | - | - | - | 1.0043 | 0.0355 | - |
| (Q) secondary contact with symmetric migration under three epoch (isolation - migration - isolation) | -3184.41 | 6380.82 | 498.9 | 5.369 | 6.5132 | - | - | - | - | 0.0803 | - | - | - | - | - | 1.6676 | 18.994 | 0.0785 |
| (L) secondary contact with asymmetric migration and population size change | -3314.64 | 6645.28 | 8242.23 | - | - | 5.8892 | 14.4296 | 0.1975 | 0.4863 | 3.1339 | 0.2216 | - | - | - | - | 0.151 | 11.1367 | - |
| (D) no migration with population size change | -3368.54 | 6749.08 | 4865.13 | - | - | 0.3808 | 0.385 | 0.8404 | 1.5644 | - | - | - | - | - | - | 0.0995 | 0.1299 | - |
| (R) secondary contact with asymmetric migration under three epoch (isolation - migration - isolation) | -3389.44 | 6792.88 | 4145.14 | 0.8947 | 0.9539 | - | - | - | - | 0.1384 | 2.1655 | - | - | - | - | 0.3917 | 0.0384 | 0.1355 |
| (A) no migration | -3730.4 | 7466.8 | 4354.57 | 0.7782 | 1.0947 | - | - | - | - | - | - | - | - | - | - | 0.3352 | - | - |
| (J) ancestral asymmetric migration | -3746.87 | 7505.74 | 4293.27 | 0.7869 | 1.112 | - | - | - | - | 28.135 | 0.7036 | - | - | - | - | 0.0305 | 0.3299 | - |
| (T) secondary contact with asymmetric migration population size change under three epoch (isolation - migration - isolation) | -3815.98 | 7649.96 | 2814.71 | - | - | 0.5965 | 5.1317 | 1.038 | 1.5606 | 23.2524 | 0.7471 | - | - | - | - | 0.9497 | 1.0888 | 0.4082 |
| (N) ancestral asymmetric migration with population size change | -3915.21 | 7846.42 | 3986.97 | - | - | 26.1339 | 1.1335 | 0.6082 | 1.1759 | 9.7451 | 0.6332 | - | - | - | - | 0.8743 | 0.2679 | - |

**Table S6.** Coordinates of the lineage-specific accentuated differentiation regions.

| chromosome | window start | window end | population pair |
|---|---|---|---|
| chr01 | 8460001 | 8470000 | BL |
| chr01 | 8470001 | 8480000 | BL |
| chr02 | 8510001 | 8520000 | BL |
| chr02 | 8520001 | 8530000 | BL |
| chr02 | 27100001 | 27110000 | BL |
| chr02 | 27110001 | 27120000 | BL |
| chr03 | 550001 | 560000 | BL |
| chr03 | 560001 | 570000 | BL |
| chr03 | 24770001 | 24780000 | BL |
| chr03 | 24780001 | 24790000 | BL |
| chr03 | 25495001 | 25505000 | BL |
| chr03 | 25505001 | 25515000 | BL |
| chr04 | 3770001 | 3780000 | BL |
| chr04 | 3780001 | 3790000 | BL |
| chr04 | 14790001 | 14800000 | BL |
| chr04 | 14800001 | 14810000 | BL |
| chr04 | 14810001 | 14820000 | BL |
| chr04 | 17070001 | 17080000 | BL |
| chr04 | 17080001 | 17090000 | BL |
| chr04 | 17180001 | 17190000 | BL |
| chr04 | 17190001 | 17200000 | BL |
| chr04 | 17820001 | 17830000 | BL |
| chr04 | 21290001 | 21300000 | BL |
| chr04 | 21300001 | 21310000 | BL |
| chr06 | 2320001 | 2330000 | BL |
| chr06 | 2330001 | 2340000 | BL |
| chr06 | 2340001 | 2350000 | BL |
| chr06 | 2350001 | 2360000 | BL |
| chr06 | 17080001 | 17090000 | BL |
| chr06 | 17090001 | 17100000 | BL |
| chr06 | 17100001 | 17110000 | BL |
| chr07 | 4095001 | 4105000 | BL |
| chr07 | 4105001 | 4115000 | BL |
| chr07 | 4115001 | 4125000 | BL |
| chr07 | 5610001 | 5620000 | BL |
| chr07 | 5620001 | 5630000 | BL |
| chr07 | 12925001 | 12935000 | BL |
| chr07 | 12935001 | 12945000 | BL |
| chr07 | 12945001 | 12955000 | BL |

| | | | |
|---|---|---|---|
| chr07 | 12965001 | 12975000 | BL |
| chr07 | 12985001 | 12995000 | BL |
| chr07 | 13015001 | 13025000 | BL |
| chr07 | 13025001 | 13035000 | BL |
| chr07 | 13515001 | 13525000 | BL |
| chr07 | 13535001 | 13545000 | BL |
| chr07 | 13545001 | 13555000 | BL |
| chr07 | 13555001 | 13565000 | BL |
| chr07 | 13655001 | 13665000 | BL |
| chr08 | 9905001 | 9915000 | BL |
| chr08 | 9915001 | 9925000 | BL |
| chr09 | 12510001 | 12520000 | BL |
| chr09 | 12520001 | 12530000 | BL |
| chr10 | 13615001 | 13625000 | BL |
| chr10 | 13625001 | 13635000 | BL |
| chr10 | 18705001 | 18715000 | BL |
| chr10 | 18715001 | 18725000 | BL |
| chr11 | 10430001 | 10440000 | BL |
| chr11 | 10440001 | 10450000 | BL |
| chr11 | 22040001 | 22050000 | BL |
| chr11 | 22050001 | 22060000 | BL |
| chr01 | 1 | 10000 | CI |
| chr01 | 10001 | 20000 | CI |
| chr01 | 1830001 | 1840000 | CI |
| chr01 | 1840001 | 1850000 | CI |
| chr01 | 3170001 | 3180000 | CI |
| chr01 | 3180001 | 3190000 | CI |
| chr01 | 5540001 | 5550000 | CI |
| chr01 | 5570001 | 5580000 | CI |
| chr01 | 5580001 | 5590000 | CI |
| chr01 | 5590001 | 5600000 | CI |
| chr01 | 5610001 | 5620000 | CI |
| chr01 | 5620001 | 5630000 | CI |
| chr01 | 7460001 | 7470000 | CI |
| chr01 | 7860001 | 7870000 | CI |
| chr01 | 7870001 | 7880000 | CI |
| chr02 | 24845001 | 24855000 | CI |
| chr02 | 24855001 | 24865000 | CI |
| chr03 | 10670001 | 10680000 | CI |
| chr03 | 10680001 | 10690000 | CI |
| chr03 | 10740001 | 10750000 | CI |
| chr03 | 10750001 | 10760000 | CI |
| chr03 | 18005001 | 18015000 | CI |

| | | | |
|---|---|---|---|
| chr03 | 18015001 | 18025000 | CI |
| chr05 | 9680001 | 9690000 | CI |
| chr05 | 9690001 | 9700000 | CI |
| chr05 | 9700001 | 9710000 | CI |
| chr05 | 10480001 | 10490000 | CI |
| chr05 | 10490001 | 10500000 | CI |
| chr05 | 15485001 | 15495000 | CI |
| chr05 | 15495001 | 15505000 | CI |
| chr06 | 14850001 | 14860000 | CI |
| chr06 | 14860001 | 14870000 | CI |
| chr06 | 20570001 | 20580000 | CI |
| chr06 | 20580001 | 20590000 | CI |
| chr07 | 2485001 | 2495000 | CI |
| chr07 | 2495001 | 2505000 | CI |
| chr07 | 2505001 | 2515000 | CI |
| chr07 | 2805001 | 2815000 | CI |
| chr07 | 2815001 | 2825000 | CI |
| chr07 | 5325001 | 5335000 | CI |
| chr07 | 5335001 | 5345000 | CI |
| chr07 | 9980001 | 9990000 | CI |
| chr07 | 9990001 | 10000000 | CI |
| chr07 | 14675001 | 14685000 | CI |
| chr07 | 14685001 | 14695000 | CI |
| chr07 | 17870001 | 17880000 | CI |
| chr07 | 17880001 | 17890000 | CI |
| chr07 | 19670001 | 19680000 | CI |
| chr07 | 19680001 | 19690000 | CI |
| chr08 | 21470001 | 21480000 | CI |
| chr08 | 21480001 | 21490000 | CI |
| chr08 | 23395001 | 23405000 | CI |
| chr08 | 24105001 | 24115000 | CI |
| chr09 | 11125001 | 11135000 | CI |
| chr09 | 11135001 | 11145000 | CI |
| chr09 | 11145001 | 11155000 | CI |
| chr10 | 290001 | 300000 | CI |
| chr10 | 300001 | 310000 | CI |
| chr10 | 310001 | 320000 | CI |
| chr10 | 13155001 | 13165000 | CI |
| chr10 | 13165001 | 13175000 | CI |
| chr10 | 13185001 | 13195000 | CI |
| chr10 | 13195001 | 13205000 | CI |
| chr10 | 13205001 | 13215000 | CI |
| chr11 | 19740001 | 19750000 | CI |

| | | | |
|---|---|---|---|
| chr11 | 19750001 | 19760000 | CI |
| chr01 | 8350001 | 8360000 | $G_{H1}N$ |
| chr01 | 8360001 | 8370000 | $G_{H1}N$ |
| chr01 | 8370001 | 8380000 | $G_{H1}N$ |
| chr01 | 8380001 | 8390000 | $G_{H1}N$ |
| chr01 | 8390001 | 8400000 | $G_{H1}N$ |
| chr01 | 8410001 | 8420000 | $G_{H1}N$ |
| chr01 | 8420001 | 8430000 | $G_{H1}N$ |
| chr01 | 16595001 | 16605000 | $G_{H1}N$ |
| chr01 | 16605001 | 16615000 | $G_{H1}N$ |
| chr02 | 22265001 | 22275000 | $G_{H1}N$ |
| chr02 | 22275001 | 22285000 | $G_{H1}N$ |
| chr03 | 12490001 | 12500000 | $G_{H1}N$ |
| chr03 | 12500001 | 12510000 | $G_{H1}N$ |
| chr04 | 7275001 | 7285000 | $G_{H1}N$ |
| chr04 | 7285001 | 7295000 | $G_{H1}N$ |
| chr04 | 13230001 | 13240000 | $G_{H1}N$ |
| chr04 | 13240001 | 13250000 | $G_{H1}N$ |
| chr04 | 17360001 | 17370000 | $G_{H1}N$ |
| chr04 | 17370001 | 17380000 | $G_{H1}N$ |
| chr05 | 3530001 | 3540000 | $G_{H1}N$ |
| chr05 | 3540001 | 3550000 | $G_{H1}N$ |
| chr06 | 4030001 | 4040000 | $G_{H1}N$ |
| chr06 | 4040001 | 4050000 | $G_{H1}N$ |
| chr06 | 4050001 | 4060000 | $G_{H1}N$ |
| chr06 | 4070001 | 4080000 | $G_{H1}N$ |
| chr06 | 4090001 | 4100000 | $G_{H1}N$ |
| chr06 | 4120001 | 4130000 | $G_{H1}N$ |
| chr06 | 4130001 | 4140000 | $G_{H1}N$ |
| chr06 | 8530001 | 8540000 | $G_{H1}N$ |
| chr06 | 8540001 | 8550000 | $G_{H1}N$ |
| chr06 | 14600001 | 14610000 | $G_{H1}N$ |
| chr06 | 14610001 | 14620000 | $G_{H1}N$ |
| chr06 | 14620001 | 14630000 | $G_{H1}N$ |
| chr06 | 14630001 | 14640000 | $G_{H1}N$ |
| chr06 | 14640001 | 14650000 | $G_{H1}N$ |
| chr06 | 14650001 | 14660000 | $G_{H1}N$ |
| chr06 | 26525001 | 26535000 | $G_{H1}N$ |

| | | | |
|---|---|---|---|
| chr06 | 26535001 | 26545000 | $G_{H1}N$ |
| chr06 | 26545001 | 26555000 | $G_{H1}N$ |
| chr08 | 2510001 | 2520000 | $G_{H1}N$ |
| chr08 | 2520001 | 2530000 | $G_{H1}N$ |
| chr08 | 2530001 | 2540000 | $G_{H1}N$ |
| chr08 | 2540001 | 2550000 | $G_{H1}N$ |
| chr08 | 4240001 | 4250000 | $G_{H1}N$ |
| chr08 | 4330001 | 4340000 | $G_{H1}N$ |
| chr08 | 4340001 | 4350000 | $G_{H1}N$ |
| chr08 | 7275001 | 7285000 | $G_{H1}N$ |
| chr08 | 7285001 | 7295000 | $G_{H1}N$ |
| chr08 | 16300001 | 16310000 | $G_{H1}N$ |
| chr08 | 16320001 | 16330000 | $G_{H1}N$ |
| chr08 | 16330001 | 16340000 | $G_{H1}N$ |
| chr08 | 16340001 | 16350000 | $G_{H1}N$ |
| chr08 | 16350001 | 16360000 | $G_{H1}N$ |
| chr08 | 16360001 | 16370000 | $G_{H1}N$ |
| chr09 | 2620001 | 2630000 | $G_{H1}N$ |
| chr09 | 2630001 | 2640000 | $G_{H1}N$ |
| chr09 | 2860001 | 2870000 | $G_{H1}N$ |
| chr09 | 2870001 | 2880000 | $G_{H1}N$ |
| chr09 | 2880001 | 2890000 | $G_{H1}N$ |
| chr09 | 12380001 | 12390000 | $G_{H1}N$ |
| chr09 | 12390001 | 12400000 | $G_{H1}N$ |
| chr10 | 14095001 | 14105000 | $G_{H1}N$ |
| chr10 | 14105001 | 14115000 | $G_{H1}N$ |
| chr10 | 14115001 | 14125000 | $G_{H1}N$ |
| chr11 | 23085001 | 23095000 | $G_{H1}N$ |
| chr11 | 23595001 | 23605000 | $G_{H1}N$ |
| chr10 | 3800001 | 3810000 | MT |
| chr10 | 3810001 | 3820000 | MT |
| chr10 | 5405001 | 5415000 | MT |
| chr10 | 5415001 | 5425000 | MT |
| chr10 | 20005001 | 20015000 | MT |
| chr10 | 20015001 | 20025000 | MT |
| chr11 | 22395001 | 22405000 | MT |
| chr11 | 22405001 | 22415000 | MT |
